# Evidence of an additional wild contributor, *Malus orientalis* Uglitzk., to the genome of cultivated apple varieties in the Caucasus and Iran

**DOI:** 10.1101/2021.03.28.437401

**Authors:** Bina Hamid, Yousefzadeh Hamed, Venon Anthony, Remoué Carine, Rousselet Agnès, Falque Matthieu, Shadab Faramarzi, Xilong Chen, Jarkyn Samanchina, David Gill, Akylai Kabaeva, Giraud Tatiana, Hossainpour Batool, Abdollahi Hamid, Gabrielyan Ivan, Nersesyan Anush, A. Cornille

## Abstract

Anthropogenic and natural divergence processes remain poorly studied in crop-wild fruit tree complexes, especially in the Caucasus, a pivotal region for plant domestication. We investigated anthropogenic and natural divergence processes in apples in the Caucasus using 26 microsatellite markers amplified in 550 wild and cultivated samples. We found two genetically distinct cultivated populations in Iran that are differentiated from *Malus domestica*, the standard cultivated apple worldwide. Coalescent-based inferences showed that these two cultivated populations originated from specific domestication events of *M. orientalis* in Iran. One of the Iranian clusters comprised both cultivated and forest trees, suggesting that either farmers use local wild apple for cultivation or that some forest trees are feral cultivars. We found evidence of substantial wild-crop and crop-crop gene flow in the Caucasus, as has been described in apple in Europe. In the Caucasus, we identified seven genetically differentiated populations of wild apple (*Malus orientalis*). Niche modeling combined with genetic diversity estimates indicated that these populations likely resulted from range changes during past glaciations. This study identifies Iran as a key region in the domestication of apple and *M. orientalis* as an additional contributor to the cultivated apple gene pool. Domestication of the apple tree therefore involved multiple origins of domestication in different geographic locations and substantial crop-wild hybridization, as found in other fruit trees. This study also highlights the impact of climate change on the natural divergence of a wild fruit tree and provides a starting point for apple conservation and breeding programs in the Caucasus.

## Introduction

Crop-wild complexes, *i.e.,* groups of crops and their associated wild species that exchange genes (Anderson, 2005), provide good models for understanding how anthropogenic and natural factors shape population divergence in the presence of gene flow. Indeed, crops are the result of a recent anthropogenic divergence process, *i.e.,* domestication, which began around 10,000 years ago, and which has often been followed by subsequent crop-wild gene flow (Besnard, Terral, & Cornille, 2018; Brandenburg et al., 2017; Chen et al., 2019; Cornille, Giraud, Smulders, Roldán-Ruiz, & Gladieux, 2014; Cornille et al., 2012; Diez et al., 2015; Flowers et al., 2019; Gaut, Díez, & Morrell, 2015). Conversely, wild species allow the study of natural divergence over a longer timescale. Indeed, wild species have often undergone shifts in their distribution following past climate changes associated with glacial periods, and range contraction has often led to population differentiation and divergence (Excoffier, Foll, & Petit, 2009; Hewitt, 1990; Hewitt, 1996; Jezkova, Olah-Hemmings, & Riddle, 2011; Petit, Bialozyt, Garnier-Géré, & Hampe, 2004; Schmitt, 2007). Understanding the evolutionary processes shaping the natural and anthropogenic divergence of crop-wild complexes is not just an academic exercise: it will also help assess the future status of wild resources. Because of the socio-economic importance of crop plants, protecting the wild relatives of crops, beyond the need for preserving biodiversity (Bacles & Jump, 2011), will allow us to manage the genetic resources for future breeding strategies in the face of global changes (*e.g.,* climate change, emerging diseases) (Bailey-Serres, Parker, Ainsworth, Oldroyd, & Schroeder, 2019; Castañeda-Álvarez et al., 2016; H. Zhang, Mittal, Leamy, Barazani, & Song, 2017).

Fruit trees present several historical and biological features that make them fascinating models for investigating anthropogenic and natural divergence with gene flow. Several fruit tree crops are found across the world and sometimes occur in sympatry with their wild relatives (Besnard, Terral, & Cornille, 2018; Cornille et al., 2019; Cornille, Giraud, Smulders, Roldán-Ruiz, & Gladieux, 2014; Delplancke et al., 2011; Liu et al., 2019; Miller & Gross, 2011). Fruit trees are also characterized by high levels of gene flow during divergence, which is to be expected considering the typical life history traits of trees (Cornille et al., 2013; Cornille, Gladieux, & Giraud, 2013; Oddou-Muratorio & Klein, 2008; Petit & Hampe, 2006). Population genetics studies of natural divergence processes associated with the last glacial maximum in Europe, North America and Asia in wind-dispersed trees (*e.g., Abies*, *Pinus*, *Fraxinus*, *Quercus*, *Betula* (Lascoux, Palmé, Cheddadi, & Latta, 2004; Petit et al., 2004)) and animal-dispersed trees (Cornille et al., 2013; George et al., 2015) showed that there were high levels of gene flow between populations and that trees had high dispersal capabilities. These studies also located single (Bai & Spitkovsky, 2010; Tian, Li, Ji, Zhang, & Luo, 2009; Zeng et al., 2011) or multiple (Qiu, Wang, Liu, Shen, & Tang, 2011; Tian et al., 2009) glacial refugia where most temperate tree species persisted during the last glacial maximum, and from which populations recolonized higher or lower latitudes during the Holocene post-glacial expansion (Giesecke, Brewer, Finsinger, Leydet, & Bradshaw, 2017). Population genetics and genomics studies also revealed the prominent role of gene flow during the anthropogenic divergence of fruit trees. Domestication of several emblematic fruit tree crops such as grape and apple occurred with substantial crop-crop and wild-crop gene flow and without a bottleneck (Arroyo-García et al., 2006; Cornille et al., 2012; Decroocq et al., 2016; Diez et al., 2015; Duan et al., 2017; Liu et al., 2019; Meyer, Duval, & Jensen, 2012; Myles et al., 2011). These studies thus revealed that the domestication of fruit trees displays different patterns from that of annuals which can be explained by the long lifespan, long juvenile phase and self-incompatibility system of trees (Fuller, 2018; Gaut et al., 2015). However, studies of natural and anthropogenic divergence processes in crop-wild fruit tree complexes are still scarce in the geographic regions that were pivotal in the divergence history of these complexes.

The Caucasus ecoregion harbors a remarkable concentration of both economically important plants and their wild relatives, in particular wheat, rye, barley and also fruit trees including walnut, apricot and apple (Asanidze, Akhalkatsi, Henk, Richards, & Volk, 2014; Gabrielian & Zohary, 2004; Vavilov, 1926, 1992; Yousefzadeh, Hosseinzadeh Colagar, Tabari, Sattarian, & Assadi, 2012). This region covers Georgia, Armenia, Azerbaijan, the North Caucasian part of the Russian Federation, the northeastern part of Turkey and the Hyrcanian Mixed Forests region in northwestern Iran (Nakhutsrishvili, Zazanashvili, Batsatsashvili, & Montalvo, 2015; Zazanashvili et al., 2020). Two Pleistocene refugia for temperate plants are recognized in this region (Bina, Yousefzadeh, Ali, & Esmailpour, 2016; Yousefzadeh et al., 2012): the Colchis refugium in the catchment basin of the Black Sea (in the Western Caucasus) and the Hyrcanian refugium at the southern edge of the Caspian Sea. Glacial refugia are known to harbor higher levels of species and genetic diversity (Hewitt, 2004), and this is the case for the Colchis and Hyrcanian refugia. Furthermore, it has been suggested that Iran, with its close proximity to Central Asia - the center of origin of emblematic fruit trees - and its historic position on the Silk Trade Routes, is a possible secondary center of domestication of apple, grape and apricot (Decroocq et al., 2016; Liang et al., 2019; Liu et al., 2019). However, inferences of the natural and anthropogenic divergence history of wild-crop fruit tree complexes in the Caucasus have been limited by the small number of samples (Decroocq et al., 2016; Liu et al., 2019) and/or genetic markers investigated so far (Amirchakhmaghi et al., 2018; Asanidze et al., 2014; Cornille et al., 2013; Gharghani et al., 2010; Myles et al., 2011; Volk & Cornille, 2019; Vouillamoz et al., 2006).

The Caucasian crab apple, *Malus orientalis* Uglitzk., is an endemic wild apple species occurring in the Caucasus. More specifically, it is found in the Western Caucasus (*i.e.*, the southern part of Russia, northern Anatolia and northwestern Turkey), Georgia, Armenia, the mountainous belt in northern Iran, the Hyrcanian Forests (Büttner, 2001; Rechinger, 1964) and the Zagros Forests in eastern and central Iran (Browicz, 1969; Rechinger, 1964) (Figure 1). This species displays high phenotypic diversity across its distribution range where it occurs as scattered individuals in natural forests or at high altitude in rocky mountains (Fischer & Schmidt, 1938; Rechinger, 1964). The Caucasian crab apple has a high resistance to pests and diseases (Büttner, 2001) and its fruit, of high quality, are variable in size (2-4 cm) and color (green to yellowish) (Cornille et al., 2014). *Malus orientalis* has been present and cultivated in Iran (formerly Persia) long before *M. domestica*, the standard cultivated apple worldwide, was introduced (Abdollahi, 2021; Curtis et al., 2005; Fallahi, Colt, Fallahi, & Chun, 2002; Fallahi, Fallahi, & Mahdavi, 2020; Gharghani et al., 2010; Ali Gharghani et al., 2009; Robert Nicholas Spengler, 2019). Translations of cuneiform scripts from Persepolis and other historical documents that are related to the Sassanid and Parthian dynasties revealed that edible apples were grown in Iran in the form of dwarf apples three centuries before Christ (BC) (Abdollahi, 2021; Curtis et al., 2005; Fallahi et al., 2002). Cultivation of *M. orientalis* in Persian gardens became more widespread after the invasion and subjugation of large portions of the Persian Empire (334 BC) by Alexander the Great (Bowe, 2004). Dwarf apples became more popular in gardens and paradise gardens (derived from the Persian term ‘pardis’) around 60 BC in Iran (Fallahi et al., 2002). Apple orchards were cultivated more widely in Iran during the Safavid period (1501- 1736), and apple cultivation has continued in Iran until now (Fallahi et al., 2002, 2020). Nowadays, the fruit of several local cultivars that are unique to Iran are harvested in the Caucasus for stewing and processed as juice and other beverages (cider, wine), jelly, syrup, jam and vinegar (Amirchakhmaghi et al., 2018; Büttner, 2001). *Malus domestica* is also currently grown in various regions of the Caucasus, but to a much lesser extent than the local cultivars (Forsline et al., 2003; Gharghani et al., 2010; Gharghani et al., 2009; Langenfeld, 1991; Schmitt, 2007). Some authors therefore suggest that the local apple cultivars from several regions of the Caucasus originated from *M. orientalis* (Forsline et al., 2003; Langenfeld, 1991; Schmitt, 2007) and not from the Central Asian wild apple, *M. sieversii,* the progenitor of *M. domestica* <u>(Cornille et al., 2019; Cornille, Giraud, Smulders, Roldán-Ruiz, & Gladieux, 2014; Duan et al., 2017, 2017; Harris, Robinson, & Juniper, 2002; Migicovsky et al., 2021)</u>, or may even be *M. orientalis* grown in gardens and orchards. Therefore, there may have been multiple independent events of domestication of the apple tree in the Caucasus, or apple domestication in the Caucasus may be in its early stage, with the cultivation of *M. orientalis* individuals in orchards. The study of the relationships between *M. orientalis*, the local Caucasian cultivars, *M. domestica* and its Central Asian progenitor, *Malus sieversii*, is therefore needed to answer the following questions: i) Are cultivated apples in the Caucasus derived from the same ancestral gene pools as *M. domestica, i.e., M. sieversii,* or are they derived from *M. orientalis*? ii) Has *M. orientalis* contributed to the local cultivated Caucasian apple germplasm through wild-to-crop introgression in the same way that *M. sylvestris,* the European crab apple, contributed to the *M. domestica* gene pool (Cornille et al., 2012; Ali Gharghani et al., 2009)?; iii) Are *M. orientalis* and/or crop-wild hybrid trees cultivated in gardens and orchards in Iran? Conversely, are there feral and/or crop-wild trees in the wild? Indeed, cultivated and wild apples can be sympatric if orchards are present near natural forests with wild apple populations. In France, the occurrence of orchards surrounding wild apple populations have directly impacted the levels of crop-to-wild introgression of the European wild apple populations (Cornille et al., 2015). However, the extent of crop-wild gene flow in apples in the Caucasus has just begun to be investigated. One study suggested that *M. orientalis* only made a minor contribution to Mediterranean *M. domestica* cultivars (Cornille et al., 2012), but lacked in-depth investigation. A population genetics study revealed low levels of crop-to-wild gene flow from *M. domestica* to *M. orientalis* in natural forests of Armenia, Turkey and Russia (Cornille et al., 2013). Population genetic diversity and structure analyses of *M. orientalis* populations from the Western Caucasus and South Caucasus identified three differentiated populations: one in Turkey, one in Armenia and one in Russia (Cornille et al., 2013). On a smaller geographic scale, an east-west genetic subdivision was found across the Hyrcanian Forests in Iran, with five main populations showing admixture (Amirchakhmaghi et al., 2018). However, we still lack a comprehensive view (beyond Armenia) of the genetic diversity and structure of *M. orientalis* to understand its natural divergence history. In addition, studying local cultivars from the Caucasus will shed light on the relationships between local *M. orientalis* apple populations, the local cultivated apple and the standard cultivated apple *M. domestica,* as well as the extent of crop-wild gene flow in apple in this region.

**Figure 1.**
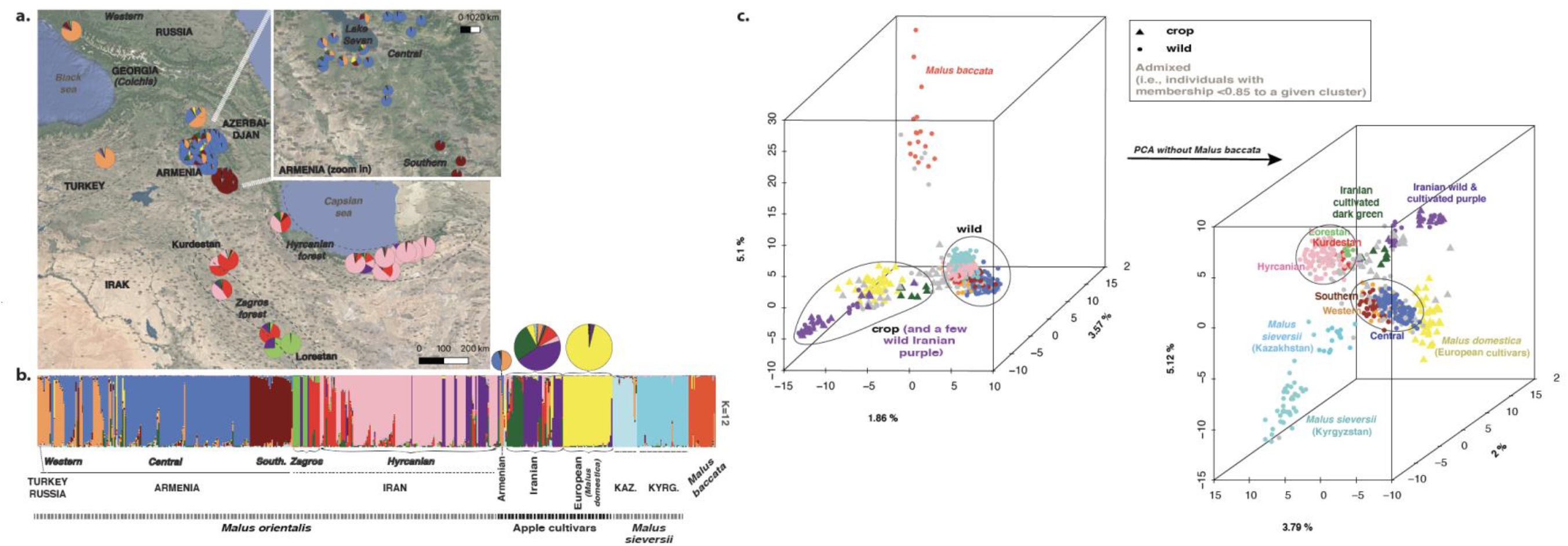
Population genetic structure and differentiation in cultivated and wild apples from the Caucasus and Iran, *Malus domestica*, *Malus sieversii* and *Malus baccata* based on 26 microsatellite markers. a. Spatial population genetic structure inferred with STRUCTURE at *K* = 12 (*N* = 550); the map represents membership proportions averaged over each geographic site for the Caucasian crabapple *M. orientalis* (*N* = 374, 43 sites across Turkey, Russia, Armenia and Iran). In the bottom right corner, the mean membership proportions for the apple cultivars from Armenia (*N* = 4), Iran (*N* = 48) and Europe (*M. domestica*, *N* = 40). The size of the pie charts is proportional to the number of samples per site. **b.** STRUCTURE bar plot (*N* = 550) at *K* = 12 showing 12 distinct genetic clusters. Each vertical line represents an individual. Colors represent the inferred ancestry from *K* ancestral genetic clusters. Sites are grouped by country for the wild apple samples (*i.e.,* Turkey, Russia, Armenia, Iran) and *M. sieversii* (*i.e.*, Kazakhstan and Kyrgyzstan), apple cultivars are grouped according to their origin: Armenia (*N* = 4), Iran (*N* = 48) and *M. domestica* (*N* = 40). Countries (Kazakhstan, Kyrgyzstan, Armenia) and/or main regions in the Caucasus (the Western Caucasus, *i.e.*, Turkey and Russia, Zagros and Hyrcanian Forests, Central and Southern Armenia) are shown on the map. Reference samples from previously published studies of each species are: *M. orientalis* from the Western Caucasus and Central and Southern Armenia, *M. domestica* (European cultivars), *M. sieversii* from Kazakhstan (Cornille et al., 2013) and *M. baccata* (Cornille et al., 2012) **c.** Principal component analysis (PCA) of 550 individuals (upper left), and after removing the outgroup *M. baccata* (lower right, *N* = 530), with the respective total variation explained by each component.

Here, we investigate anthropogenic and natural divergence processes in apples from the Caucasus and the extent of gene flow during divergence. A total of 550 apple trees, comprising local cultivated and wild apples from the Caucasus, *M. domestica* apple cultivars and *M. sieversii,* were genotyped using 26 microsatellite markers. In addition, the Siberian wild apple *Malus baccata* was used as an outgroup in certain analyses. From the analysis of this comprehensive genetic dataset, combined with ecological niche modeling approaches, we addressed the following questions: 1) What is the population genetic structure among Caucasian wild and cultivated apples, *M. domestica* and *M. sieversii,* and what are their genetic relationships and levels of genetic diversity? 2) What is the extent of crop-crop, crop-wild and wild-wild gene flow in apples in the Caucasus? Are there feral or crop-wild hybrid trees in the wild or, conversely, do humans cultivate wild or hybrid trees? 3) Is *M. orientalis* an additional contributor to the domestication history of cultivated apple? 4) Did *M. orientalis* experience past range contraction and expansion associated with the last glacial maximum?

## Materials and methods

### Sampling, DNA extraction and microsatellite genotyping

Microsatellite genotyping data for *M. orientalis*, *M. sieversii* from Kazakhstan, *M. domestica* and *M. baccata* were from previously published studies, including: 207 *M. orientalis* individuals from Turkey, Armenia and Russia (23 sites, Tables S1 and S2, (Cornille et al., 2013; Cornille et al., 2012)), four apple cultivars from Armenia, 40 “pure” European cultivated *M. domestica* individuals (*i.e.*, not introgressed by *M. sylvestris*) (Tables S1 and S2, (Cornille et al., 2013, 2012)), 20 *M. sieversii* individuals from Kazakhstan (Cornille et al., 2013) and 22 *M. baccata* individuals from Russia (Cornille et al., 2012). We collected 257 new samples in 2017 in Iran and in 2018 in Kyrgyzstan (*M. sieversii*) for this study: 167 *M. orientalis* individuals from the Hyrcanian Forests and the Zagros region in Iran (Table S1), 48 local Iranian apple cultivars from the *Seed and Plant Improvement Institute* (Karaj, Iran) (Table S1) and 42 *M. sieversii* individuals from Kyrgyzstan. Note that for 18 of the 48 local Iranian samples, we measured fruit size (Table S1). Collections meet the requirements of the recently enacted Nagoya protocol on access to genetic resources and the fair and equitable sharing of benefits. Thus, a total of 550 individuals were analyzed, comprising 374 *M. orientalis*, 48 Iranian and four Armenian apple cultivars, 40 European apple cultivars belonging to *M. domestica,* 62 *M. sieversii* (from Kyrgyzstan and Kazakhstan) and 22 *M. baccata* individuals (details are provided in Table S1).

DNA from the new samples (*N* = 257) was extracted from dried leaves with the NucleoSpin plant II DNA extraction kit (Macherey & Nagel, Düren, Germany®) following the manufacturer’s instructions. Multiplex microsatellite PCR amplifications were performed with a multiplex PCR kit (Qiagen Inc.®) for 26 microsatellite markers as previously described (Cornille et al., 2012; Patocchi, Frei, Frey, & Kellerhals, 2009). Note that on each DNA plate, we included three controls, *i.e.,* one sample of *M. orientalis,* one of *M. sieversii* and one of *M. domestica* for which data were already available (Cornille et al., 2013). Genotypes of the controls for each of the 26 microsatellite markers were compared with the 2013 dataset. We retained only multilocus genotypes for which < 20 % of the data was missing. The suitability of these markers for population genetics analyses has been demonstrated in previous studies (Cornille et al., 2013; Cornille, Gladieux, & Giraud, 2013; Cornille et al., 2012).

### Bayesian inferences of population structure, genetic differentiation among wild and cultivated apples, and genetic diversity estimates

We investigated the population structure of wild and cultivated apples with the individual-based Bayesian clustering methods implemented in STRUCTURE 2.3.3 (Pritchard, Stephens, & Donnelly, 2000). STRUCTURE uses Markov chain Monte Carlo (MCMC) simulations to infer the proportion of ancestry of genotypes from *K* distinct clusters. The underlying algorithms attempt to minimize deviations from Hardy–Weinberg within clusters and linkage disequilibrium among loci. We ran STRUCTURE from *K* = 1 to *K* = 15. Based on 10 repeated runs of MCMC sampling from 500,000 iterations after a burn-in of 50,000 steps, we determined the amount of additional information explained by increasing *K* using the *ΔK* statistic (Evanno, Regnaut, & Goudet, 2005) as implemented in the online post-processing software Structure Harvester (Earl, 2012). However, *ΔK* provides statistical support for the strongest but not the finest population structure (Puechmaille, 2016). Natural populations can display hierarchical genetic structure, with fine-scale population structure. It has been shown that interpreting *K* values below the one corresponding to the finest substructure can lead to misinterpretations (Puechmaille, 2016). Indeed, simulations have shown that STRUCTURE is not reliable for identifying groups that are not at Hardy-Weinberg equilibrium, *i.e.*, when the *K* is too low (Cullingham et al., 2020; Kalinowski, 2011; Puechmaille, 2016). For instance, when the number of clusters is suboptimal, STRUCTURE does not always group the genetically closest clusters into the same cluster. The problem seems to be caused when the program is forced to cluster individuals into an inappropriately small number of clusters (Cullingham et al., 2020; Kalinowski, 2011; Puechmaille, 2016). A method for identifying the best *K* value is to look at the bar plots and choose the *K* value for which all clusters have well assigned individuals, and where additional clusters at higher *K* values do not have well assigned individuals (indicating that we have reached the highest *K* value for which new genuine clusters could be delimited). The *K* value we considered therefore corresponded to the finest one, which can be higher than the *K* value of the strongest population structure level identified by *ΔK*.

We ran STRUCTURE for the whole dataset (*N* = 550) to investigate the population genetic structure among the Caucasian crab apple *M. orientalis*, the Caucasian and Iranian cultivated apples, *M. sieversii*, *M. domestica* and *M. baccata*. We further explored the genetic variation and differentiation among the genetic groups detected with STRUCTURE using three different methods. First, we ran a principal component analysis (PCA) for all individuals with the *dudi.pca* function from the *adegenet* R package (Jombart & Ahmed, 2011). For the PCA, individuals that were assigned to a given cluster with a membership coefficient ≥ 0.85 were color-coded according to the respective color of each cluster, and admixed individuals (*i.e.,* individuals with a membership coefficient to any given cluster < 0.85) were colored in gray. We chose this threshold based on the distribution of the maximum membership coefficients inferred with STRUCTURE (see results). The angle used to draw the PCA was 380 degrees for a better visualization of the data. Second, we generated a neighbor-net with Splitstree v4 (Huson, 1998; Huson & Scornavacca, 2012), using the PCA color code. Third, we explored the relationships among populations identified with STRUCTURE (*i.e.,* clusters of individuals with a membership coefficient ≥ 0.85 to a given cluster) with a neighbor joining (NJ) tree (Huson, 1998; Huson & Scornavacca, 2012). The NJ tree and the neighbor-net tree were built using Nei’s standard genetic distance (Nei, 1987) computed among individuals or populations with the Populations software v1.2.31 (https://bioinformatics.org/populations/).

We computed descriptive population genetic estimates for each population (*i.e.,* each cluster inferred with STRUCTURE, excluding admixed individuals with a membership coefficient < 0.85). We calculated allelic richness (*A_R_*) and private allelic richness (*A_P_*) with ADZE (Szpiech, Jakobsson, & Rosenberg, 2008) using standardized sample sizes of *N_ADZE_* = 7 (one individual x two chromosomes), corresponding to the minimal number of observations across populations. Heterozygosity (expected and observed), Weir and Cockerham *F-*statistics and deviations from Hardy–Weinberg equilibrium were calculated with Genepop v4.2 (Raymond & Rousset, 1995; Rousset, 2008).

### Identification of crop-wild hybrids in the Caucasus and historical wild-wild gene flow in the Caucasian crabapple, *M. orientalis*

To assess the extent of crop-wild gene flow in the Caucasus, we removed *M. sieversii* and *M. baccata* from the dataset (resulting in a dataset with *N* = 466, Table S2) and ran STRUCTURE with the same parameters as above. We defined hybrids resulting from crop-to-wild introgression as *M. orientalis* trees assigned to the *M. domestica* or the Iranian or Armenian cultivated gene pools with a membership coefficient **>** 0.10. We defined hybrids resulting from wild-to-crop introgression as cultivars assigned to any of the wild gene pools with a membership coefficient **>** 0.10. We chose this threshold based on the distribution of the maximum membership coefficients inferred with STRUCTURE (see results).

After removing crop-wild hybrids, we estimated the extent of historical wild-wild gene flow in *M. orientalis* in the Caucasus using two methods. First, we tested whether there was a significant isolation-by-distance (IBD) pattern. We computed the correlation between *F_ST_/(1-F_ST_)* and the natural algorithm of geographic distance with SPAGeDI 1.5 (Hardy & Vekemans, 2002). Second, for each population, we computed the Nason’s kinship coefficient *F_ij_* between pairs of individuals *i* and *j* (Loiselle, Sork, Nason, & Graham, 1995)) with SPAGeDI 1.5 (Hardy & Vekemans, 2002), and regressed *F_ij_* against the natural logarithm of geographic distance, *ln(d_ij_),* to obtain the regression slope *b*. We permuted the spatial position of individuals 9,999 times to test whether there was a significant spatial genetic structure between sites. We then calculated the *Sp* statistic, defined as *Sp = -bLd/(1-F_N_)*, where *F_N_* is the mean *F_ij_* between neighboring individuals (Vekemans & Hardy, 2004), and *-bLd* is the regression slope of *F_ij_* against *ln(d_ij_)*. A low *Sp* implies low spatial population structure, which suggests high historical gene flow and/or high effective population size.

We investigated the directionality of crop-wild, wild-wild and crop-crop gene flow (*i.e.*, immigration rate *m_ji_*, the proportion of immigrants in population*i* that arrive from population *j*) over the past few generations with a Bayesian assignment method implemented in BAYESASS v3.0 (Wilson & Rannala, 2003). As recommended by Wilson and Rannala (Wilson & Rannala, 2003), we run 3×10^6^ iterations, sampled every 2,000 iterations with a discarded burn-in of 10^6^ iterations, delta values were adjusted to 0.12 to ensure that chain swapping occurred in approximately 50% of the total iterations. We repeated five times the analysis with different random number seeds to evaluate consistency of results.

### Approximate Bayesian computation to reconstruct the domestication history of apple

We used approximate Bayesian computation to test whether Iranian cultivated apples (see results below, we removed Armenian cultivated apples as they were represented by only four samples) were derived from the same domestication event as *M. domestica*. We used the newly developed ABC method based on a machine learning tool named “random forest” (ABC-RF) to perform model selection and parameter estimates (Estoup et al., 2018; Pudlo et al., 2016; Raynal et al., 2019). In brief, this method creates a “forest” of bootstrapped decision trees that ranks scenarios based on the summary statistics of the datasets. Some simulations are not used to build the trees and can thus be used to cross-validate the analysis by computing a “prior error rate”. This approach allows the comparison of complex demographic models (Pudlo et al., 2016) by comparing groups of scenarios with a specific type of evolutionary event with other groups with different types of evolutionary events instead of considering all scenarios separately (Estoup et al., 2018).

Using the ABC-RF framework, we compared different scenarios of domestication of Iranian cultivars, *i.e.*, with an origin from *M. domestica*, *M. sieversii* from Kazakhstan or *M. orientalis*. Populations were defined as the clusters detected with STRUCTURE with the whole dataset (*N* = 550), removing putative hybrid individuals (*i.e.*, individuals with a membership coefficient < 0.85 to any given cluster), and excluding *M. baccata* and Armenian cultivars (see Results part). We removed putative hybrids in order to retrace the divergence history and historical gene flow among populations; more recent admixture events being detectable directly from the STRUCTURE bar plots. We assumed bidirectional gene flow among wild and cultivated apple populations. The model parameters used were: the divergence time between *X* and *Y* populations (*T_X-Y_*), the effective population size of population *X* (*N_E-X_*) and the migration rate per generation between *X* and *Y* populations (*m_X-Y_*). Prior values for divergence time were drawn from the log-uniform distribution bounded between the distributions used in the approximate Bayesian computations and are given in Table S3.

For all models, microsatellite datasets were simulated for 14 out of the 26 markers that had perfect repeats (Ch01h01, Ch01h10, Ch02c06, Ch02d08, Ch05f06, Ch01f02, Hi02c07, Ch02c09, Ch03d07, Ch04c07, Ch02b03b, MS06g03, Ch04e03, Ch02g01 (Cornille et al., 2013; Cornille et al., 2012; Patocchi, Fernàndez-Fernàndez, et al., 2009). We checked that the population structure inferred with 14 microsatellite markers did not differ significantly from the inferences obtained with 26 SSR markers (data not shown). We assumed a generalized stepwise model of microsatellite evolution (Slatkin, 1995). Mutation rates were allowed to vary across loci, with locus-specific mutation rates where *μ* is the mutation rate per generation, with a log-uniform prior distribution for *μ* (10^e-4^, 10^e-3^) (Table S3).

We used ABCtoolbox (Wegmann, Leuenberger, Neuenschwander, & Excoffier, 2010) with fastsimcoal 2.5 (Excoffier & Foll, 2011) to simulate datasets, using model parameters drawn from prior distributions (Table S3). We performed 8,000 simulations per scenario. For each simulation, we calculated three summary statistics per population with arlsumstats v 3.5 (Excoffier and Lischer 2010): *H*, the mean heterozygosity across loci, *GW,* the mean Garza-Williamson statistic over populations (Garza & Williamson, 2001) and the pairwise *F_ST_* between populations (Weir & Cockerham, 1984).

We used the abcrf v.1.7.0 R statistical package (Pudlo et al., 2016) to carry out the ABC-RF analysis. This analysis provides a classification vote that represents the number of times a scenario is selected as the best one among *n* trees in the constructed random forest. For each ABC step, we selected the scenario, or the group of scenarios, with the highest number of classification votes as the best scenario, or best group of scenarios, among a total of 500 classification trees (Breiman, 2001). We computed the posterior probabilities and prior error rates (*i.e.*, the probability of choosing a wrong group of scenarios when drawing model index and parameter values from the priors of the best scenario) over 10 replicate analyses (Estoup et al., 2018) for each ABC step. We also checked visually that the simulated models were compatible with the observed dataset by projecting the simulated and the observed datasets onto the first two linear discriminant analysis (LDA) axes (Pudlo et al., 2016), and by checking that the observed dataset fell within the clouds of simulated datasets. We then calculated parameter inferences using the final selected model. The median and 90% confidence interval (CI, 5%-95%) are given for each model parameter estimate. Note that the ABC-RF approach includes the model checking step that was performed *a posteriori* in previous ABC methods.

### Spatial pattern of genetic diversity in the Caucasian crab apple

We investigated spatial patterns of diversity in “pure” *M. orientalis*. To this aim, we excluded the crop-to-wild hybrids detected in the second STRUCTURE analysis (*i.e.,* excluding *M. baccata* and *M. sieversii*), as well as *M. domestica* and the Iranian and Armenian cultivars. Spatial patterns of genetic diversity in “pure” *M. orientalis* were visualized by mapping the variation across space (*A_R_*) at 36 sites (*i.e.,* geographic locations for which at least five individuals were successfully genotyped for each marker, Table S2) with the geometry-based inverse distance weighted interpolation in QGIS (Quantum GIS, GRASS, SAGA GIS). We calculated allelic richness (*A_R_*) and private allelic richness (*A_P_*) per site with ADZE (Szpiech et al., 2008) using standardized sample sizes of *N_ADZE_* = 6 (one individual x two chromosomes), corresponding to the minimal number of observations across sites.

### Species distribution modeling

The BIOMOD2 R package (Thuiller, Georges, Engler & Breiner, 2016) was used to project past and present distributions of *M. orientalis* following the species distribution modeling methods of Leroy *et al.* (2014). A set of 19 bioclimatic variables from WorldClim.org was used in addition to monthly temperature and precipitation values. Climate data were obtained for past conditions from the last glacial maximum and for the current period between 1960 and 1990. The climate projection at the 2.5-minute spatial resolution from the CCSM4 global climate model was used (https://www.worldclim.org/data/worldclim21.html#), as we previously showed that it was the most accurate for apple trees (Cornille et al., 2013). Past and present distributions were projected using three modeling algorithms: a generalized linear model (GLM), a generalized additive model (GAM) and artificial neural networks (ANN).

The location of 339 “pure” *M. orientalis* trees (*i.e.,* individuals assigned to a wild apple gene pool with a membership coefficient > 0.9, see results from the second STRUCTURE analysis) provided the longitude and latitude coordinates. Duplicate data points were removed, resulting in 57 presence points for *M. orientalis* (Table S4). We did not have absence data so we randomly selected pseudo-absences to serve as “absence” points for the model, and weighted presence and absence points equally as per Barbet-Massin *et al.* (2012). Models were calibrated using the set of bioclimatic variables and model evaluation was calculated with Jaccard’s indices. Ensemble model forecasting was completed by pulling the average trend of the three modeling algorithms and retaining only the uncorrelated bioclimatic variables with a Pearson correlation threshold greater than 0.75 (Table S5). The model was run again using only variables with high predictive power.

## Results

### Clear population genetic structure among Caucasian wild and cultivated apples, *M. domestica* and *M. sieversii*

The *ΔK* statistic indicated that the strongest level of population subdivision was at *K* = 3 (Figure S1 a, b). However, further genetic subdivisions were observed for *K* > 3, with well delimited and biologically meaningful clusters. We therefore visually examined the bar plots and chose the *K* value at which all clusters had fully assigned individuals, indicating the finest level of genetic subdivision. At *K* = 12, STRUCTURE identified twelve well-delimited clusters (Figures 1 a, b and S2) corresponding to species and/or geographic regions (Figure 1). We therefore considered these twelve clusters as the most relevant genetic structure.

Among these twelve clusters were two distinct genetic clusters of *M. sieversii*, one from Kazakhstan and one from Kyrgyzstan (in two shades of light blue, respectively, Figure 1), and a specific genetic cluster of *M. baccata* (orange red). We identified seven distinct genetic groups of *M. orientalis*: a genetic group from the Western Caucasus (Russia, Turkey and northwestern Armenia; orange), a central Armenian group (blue), a southern Armenian group (brown), and four genetic groups in Iran corresponding to two gene pools spread across the Zagros Forests (including samples from the Lorestan province in light green and from the Kurdestan province in red), and two gene pools (pink and purple) spread across the Hyrcanian Forests (Figure 1 a, b).

The *M. domestica* apple cultivars formed a specific genetic group (yellow) that was well separated from the wild *M. orientalis* and the Iranian and Armenian cultivars. The Iranian apple cultivars formed two gene pools: one that included only cultivars (dark green), and another (purple) that included cultivated trees and wild individuals from the Hyrcanian Forests. We also detected Iranian cultivated trees that were highly admixed between the Iranian cultivated dark green cluster, the Iranian purple cluster, the *M. domestica* yellow cluster and with two other clusters (red and orange); the latter two included several wild *M. orientalis* individuals from the Zagros Forests in the Kurdestan province in Iran and the Western Caucasus, respectively (Figures 1 and S2). The four Armenian cultivated apples fell within the blue and orange clusters, which also included *M. orientalis* trees.

We assigned individuals with a membership coefficient > 0.85 to a given cluster to the corresponding population (Figure S3) to assess the genetic variation among wild and cultivated apples. The PCA (Figure 1c) showed that *M. baccata* was highly differentiated from the other genetic groups (Figure 1c, upper left). The European (yellow) and the two Iranian cultivated genetic clusters (green and purple) formed well differentiated gene pools (Figure 1c, lower right). *Malus orientalis* from the Western Caucasus (orange) and Central and Southern Armenia (brown and blue) were closer to each other than to the Iranian wild apples (light green, red and pink), which clustered together. *Malus sieversii* from Kazakhstan was closer to the Armenian and Iranian wild apple than *M. sieversii* from Kyrgyzstan.

STRUCTURE analysis and PCA thus revealed insights into the history of apples in the Caucasus. First, cultivated apples in Iran (dark green and purple) may have resulted from additional domestication events, distinct from that of *M. domestica*, as their respective clusters are not the most closely related in the PCA. The close genetic relationships between the wild and cultivated Iranian clusters (Figure 1 and 2a) indicates that these additional domestication events could have involved *M. orientalis* in Iran; alternatively, cultivated apples in Iran could be derived from *M. sieversii* or *M. domestica* with subsequent local gene flow from *M. orientalis*. Second, the full membership of a substantial number of cultivated trees to genetic clusters of Iranian and Armenian *M. orientalis* suggests that wild trees are grown in orchards for consumption without any strong domestication process and/or that feral individuals occur (*e.g.,* the purple genetic cluster may represent a cultivated group that is also found in the wild as feral). Third, the high level of admixture in several Iranian cultivated apple trees with wild *M. orientalis* gene pools indicates substantial wild-crop gene flow. Fourth, the spatial population structure of *M. orientalis* in the Caucasus may result from past range contraction and expansion associated with the last glacial maximum. We tested these hypotheses below. First, we investigated the genetic diversity and differentiation among Caucasian wild and cultivated apples, *M. domestica* and *M. sieversii*. Second, we estimated the extent of crop-wild and wild-wild gene flow in apples in the Caucasus (*i.e.*, excluding *M. baccata* and *M. sieversii*). Third, we inferred the anthropogenic divergence history of the Iranian cultivated apple using a coalescent-based inference method for assessing the probability of different domestication scenarios. Fourth, we investigated both spatial genetic distribution of genetic diversity and modeled the past and present climatic niches of *M. orientalis* to test whether this species underwent past contraction and expansion in the Caucasus during past glaciations.

**Figure 2.**
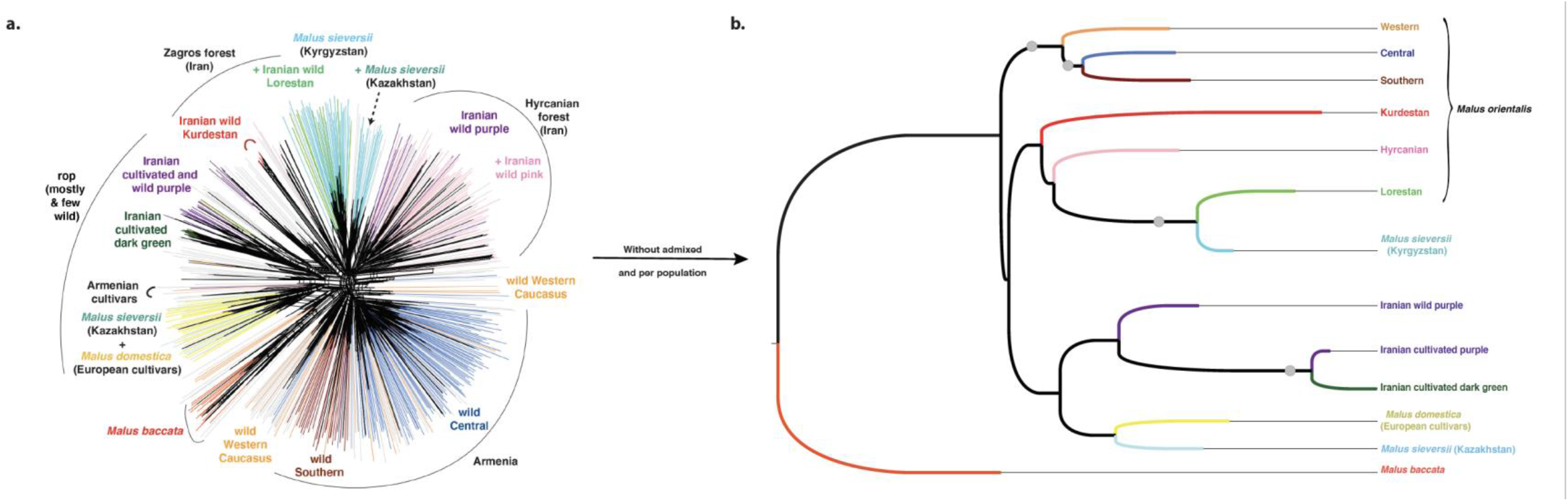
Genetic differentiation among cultivated and wild apples from the Caucasus, *Malus domestica*, *Malus sieversii* and *Malus baccata* based on 26 microsatellite markers. a. Neighbor-net representing the genetic relationships among wild and cultivated individuals inferred with STRUCTURE at *K* = 12. Colors correspond to the genetic groups inferred with STRUCTURE at *K* = 12 and admixed samples are in grey. b. Neighbor-joining tree representing the distance among the twelve populations detected with STRUCTURE at *K* = 12, excluding admixed individuals (*i.e.*, individuals with a membership coefficient < 0.85 to any given cluster), and rooted with *M. baccata*. Each branch is coloured according to the population inferred with STRUCTURE at *K* = 12, nodes with a grey circle represent bootstrap values > 80%.

### Genetic diversity and differentiation among wild and cultivated apples

The neighbor-net tree (Figure 2a) and the NJ tree (Figure 2b) confirmed that the apple cultivars (*M. domestica* and the purple and dark green Iranian cultivar clusters) were distinct from the wild populations (with the exception of the wild Hyrcanian purple group, see below). Note that we excluded admixed individuals (*i.e.*, individuals assigned with a membership coefficient < 0.85 to a given cluster) from the NJ analysis to better assess the genetic relationships among pure cultivated and wild populations.

Individuals of *M. domestica* and *M. sieversii* from Kazakhstan were intermingled (Figure 2a) and the genetic clusters of these species were sister groups (Figure 2b). Neither of the two distinct Iranian cultivated gene pools (green and purple) was sister to *M. domestica*, supporting the view that specific domestication events occurred in Iran. The two cultivated apple populations from Iran were genetically highly differentiated (Table S6), were sister groups (Figure 2) and had lower levels of genetic diversity and fewer private alleles than *M. domestica* (*P* < 0.01, Tables 1 and S7); the purple gene pool displayed the lowest level of genetic diversity and the least number of private alleles. The close relationship between trees sampled in the Hyrcanian Forests and cultivars from the purple gene pool (Figures 1 and 2), together with the lower levels of genetic diversity in these populations, suggest that the trees sampled in the Hyrcanian Forests assigned to the purple genetic group may be feral. Alternatively, the cultivated trees from the purple gene pool may represent the first step of apple domestication in Iran, *i.e.,* wild trees cultivated by humans.

**Table 1.**
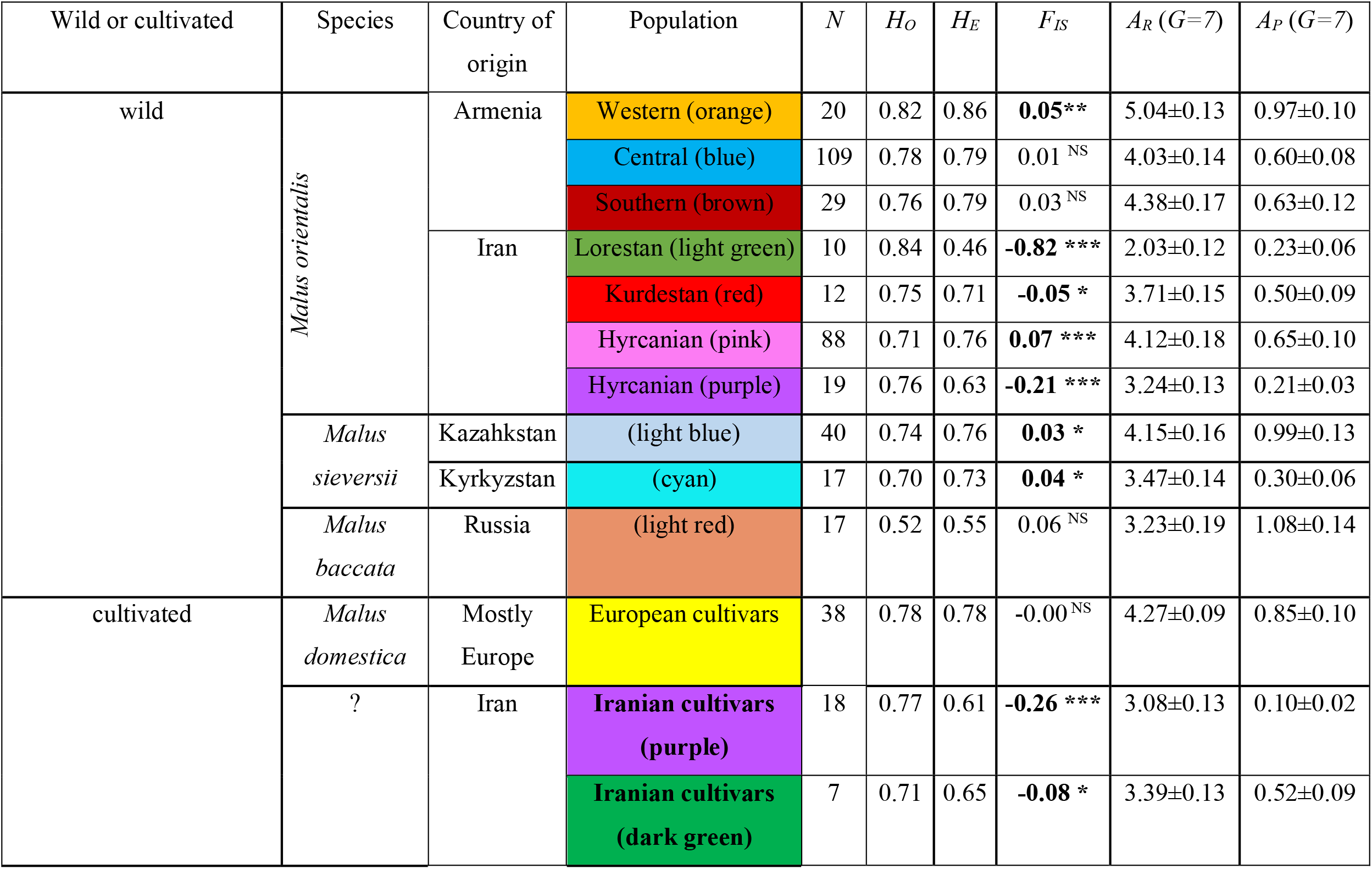
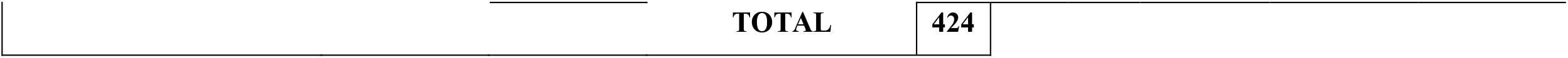
Genetic diversity estimates for wild and cultivated apple populations detected with STRUCTURE at *K* = 12. (*N* = 424, *i.e.,* individuals with a membership coefficient < 0.85 to any given cluster were excluded from the analysis). Note that the purple cluster was split between cultivated and wild samples. Thus, samples were partitioned into 13 populations, including 10 wild and three cultivated apple populations.

| Wild or cultivated | Species | Country of origin | Population | $N$ | $H_O$ | $H_E$ | $F_{IS}$ | $A_R (G=7)$ | $A_P (G=7)$ |
| --- | --- | --- | --- | --- | --- | --- | --- | --- | --- |
| wild | *Malus orientalis* | Armenia | Western (orange) | 20 | 0.82 | 0.86 | **0.05\*\*** | 5.04±0.13 | 0.97±0.10 |
|  |  |  | Central (blue) | 109 | 0.78 | 0.79 | 0.01<sup>NS</sup> | 4.03±0.14 | 0.60±0.08 |
|  |  |  | Southern (brown) | 29 | 0.76 | 0.79 | 0.03<sup>NS</sup> | 4.38±0.17 | 0.63±0.12 |
|  |  | Iran | Lorestan (light green) | 10 | 0.84 | 0.46 | **-0.82 \*\*\*** | 2.03±0.12 | 0.23±0.06 |
|  |  |  | Kurdestan (red) | 12 | 0.75 | 0.71 | **-0.05 \*** | 3.71±0.15 | 0.50±0.09 |
|  |  |  | Hyrceanian (pink) | 88 | 0.71 | 0.76 | **0.07 \*\*\*** | 4.12±0.18 | 0.65±0.10 |
|  |  |  | Hyrceanian (purple) | 19 | 0.76 | 0.63 | **-0.21 \*\*\*** | 3.24±0.13 | 0.21±0.03 |
|  | *Malus sieversii* | Kazakhstan | (light blue) | 40 | 0.74 | 0.76 | **0.03 \*** | 4.15±0.16 | 0.99±0.13 |
|  |  | Kyrkyzstan | (cyan) | 17 | 0.70 | 0.73 | **0.04 \*** | 3.47±0.14 | 0.30±0.06 |
|  | *Malus baccata* | Russia | (light red) | 17 | 0.52 | 0.55 | 0.06<sup>NS</sup> | 3.23±0.19 | 1.08±0.14 |
| cultivated | *Malus domestica* | Mostly Europe | European cultivars | 38 | 0.78 | 0.78 | -0.00<sup>NS</sup> | 4.27±0.09 | 0.85±0.10 |
|  | ? | Iran | Iranian cultivars (purple) | 18 | 0.77 | 0.61 | **-0.26 \*\*\*** | 3.08±0.13 | 0.10±0.02 |
|  |  |  | Iranian cultivars (dark green) | 7 | 0.71 | 0.65 | **-0.08 \*** | 3.39±0.13 | 0.52±0.09 |

|  |  |  |
| --- | --- | --- |
|  | **TOTAL** | **424** |

*Malus orientalis* and *M. sieversii* did not form a monophyletic group (Figure 2b). Wild *M. orientalis* populations from the Western Caucasus (orange) and from central (blue) and southern (brown) Armenia grouped together, the latter two being sister groups. The *M. orientalis* population from the Lorestan province (Iran; light green) was intermingled with the *M. sieversii* population from Kyrgyzstan (cyan), (Figure 2a); when considered as separate populations (*i.e.*, excluding admixed individuals Figure 2b), the two populations formed sister groups (Figure 2b). Although the *M. orientalis* Hyrcanian (pink and purple) populations were intermingled (with the exception of a few wild purple individuals clustering with the cultivated purple population) in the neighbor-net, the NJ tree indicated that the wild purple population was closer to the cultivated purple population. Some *M. sieversii* trees from Kazakhstan were intermingled with *M. sieversii* from Kyrgyzstan and clustered with the wild Iranian populations. *Malus sieversii* from Kazakhstan formed a distinct group, placed as sister group to *M. domestica* in both neighbor-net and NJ tree (Figure 2). The level of allelic richness was significantly lower in the wild apple populations from the Zagros Forests (*i.e.*, Lorestan (light green) and Kurdestan (red)) than in the other wild populations (Tables 1 and S7).

### Substantial crop-wild, crop-crop and wild-wild gene flow in apples in the Caucasus

The second STRUCTURE analysis, focused on the Caucasus, revealed the same genetic clustering for wild apples and *M. domestica* at *K* = 9 (Figure S4) as in the previous analysis (*K* = 12) (Figures 1 and 2). At *K* = 9, 150 apple genotypes could be considered hybrids (*i.e.,* individuals assigned to a gene pool with a membership coefficient < 0.9, this cut-off being chosen on the basis of the distribution of the cumulated membership coefficients for each individual at *K* = 9, Figure S5); these 150 hybrids represented 32% of the total dataset (Table 2). The Iranian cultivars had the highest proportion of hybrids (67%), mostly admixed with the wild and cultivated gene pools from Iran, but also with the *M. domestica* gene pool. Hybrids of the wild Armenian apple were mostly an admixture of the wild Armenian gene pools (*i.e.,* Western, Central and Southern), suggesting local gene flow between crop and wild populations.

**Table 2.**
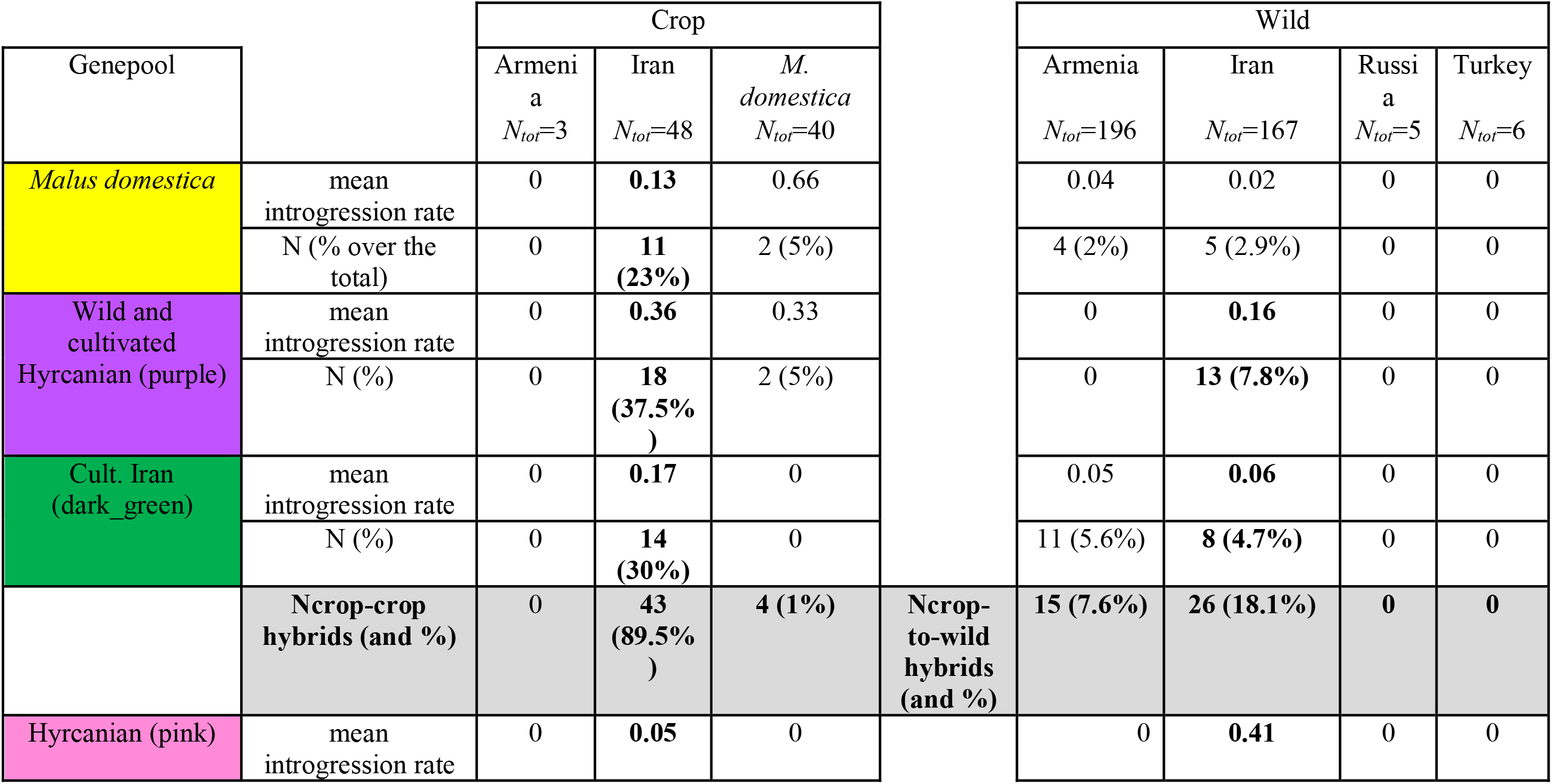
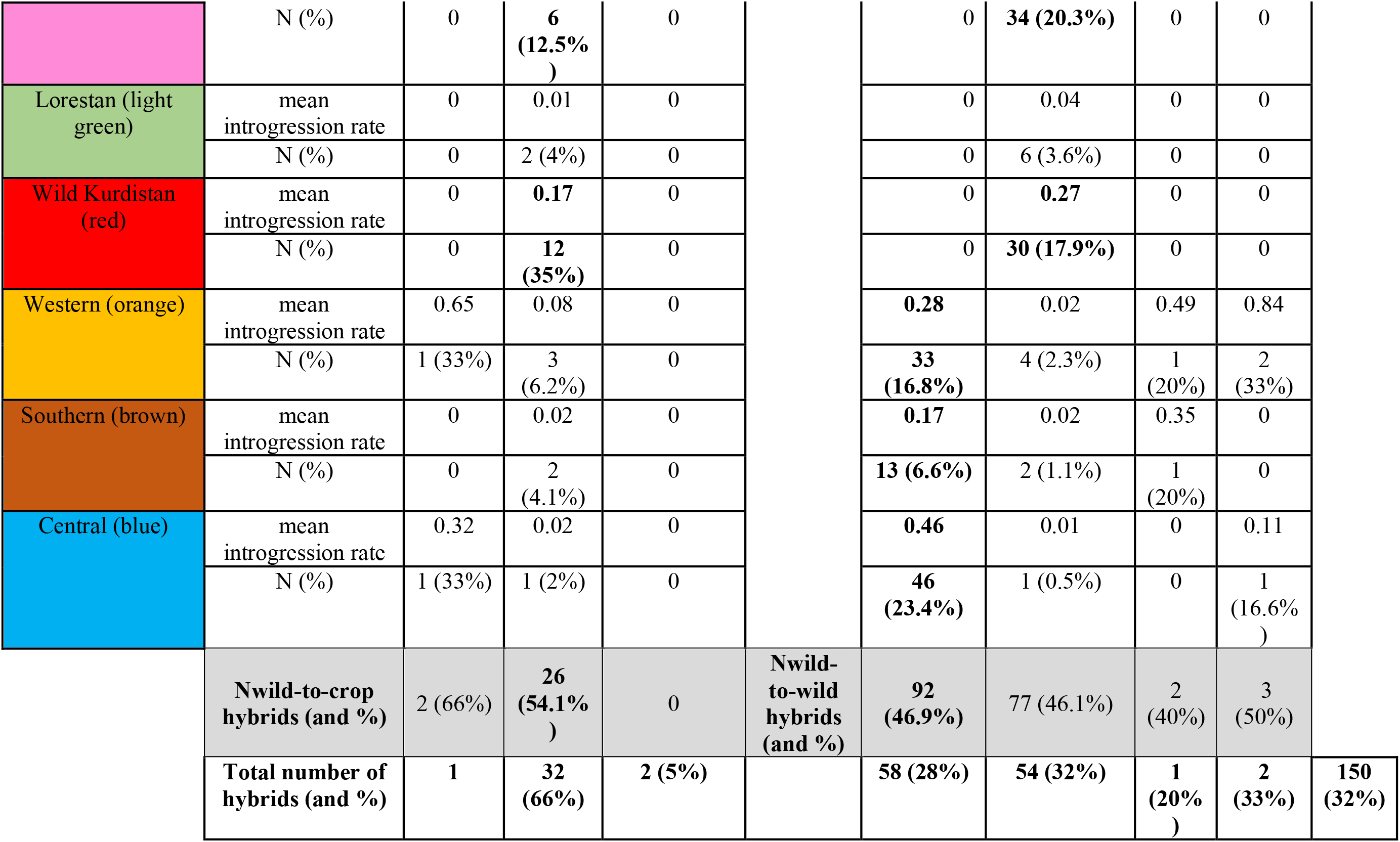
Distribution of hybrids (*i.e.*, individuals with a membership coefficient < 0.90 to any given genetic cluster, as inferred with STRUCTURE for *K* = 9) in cultivated and wild apple in the Caucasus (*N* = 466, 26 microsatellite markers). For each group (cultivated or wild, from different regions), *N_tot_* is the total number of samples in each group, *N* is the number of hybrids assigned to each gene pool and % is the respective percentage over the total number of samples from each group, the mean introgression rate is the mean membership coefficient to this gene pool. Note that some admixed trees were assigned to several gene pools with a membership coefficient < 0.90; the total number of hybrids associated with each cluster (TOTAL) is given on the last line of the table. We also showed the distribution of crop-crop, crop-to-wild, wild-to-crop and wild-to-wild hybrids.

| Genepool |  | Crop |  |  |  | Wild |  |  |  |
| --- | --- | --- | --- | --- | --- | --- | --- | --- | --- |
| | | Armeni<br>a<br>$N_{tot}=3$ | Iran<br>$N_{tot}=48$ | *M.*<br>*domestica*<br>$N_{tot}=40$ | | Armenia<br>$N_{tot}=196$ | Iran<br>$N_{tot}=167$ | Russi<br>a<br>$N_{tot}=5$ | Turkey<br>$N_{tot}=6$ |
| *Malus domestica* | mean<br>introgression rate | 0 | **0.13** | 0.66 |  | 0.04 | 0.02 | 0 | 0 |
|  | N (%) over the<br>total) | 0 | **11**<br>**(23%)** | 2 (5%) |  | 4 (2%) | 5 (2.9%) | 0 | 0 |
| Wild and<br>cultivated<br>Hyrcanian (purple) | mean<br>introgression rate | 0 | **0.36** | 0.33 |  | 0 | **0.16** | 0 | 0 |
|  | N (%) | 0 | **18**<br>**(37.5%**<br>**)** | 2 (5%) |  | 0 | **13 (7.8%)** | 0 | 0 |
| Cult. Iran<br>(dark_green) | mean<br>introgression rate | 0 | **0.17** | 0 |  | 0.05 | **0.06** | 0 | 0 |
|  | N (%) | 0 | **14**<br>**(30%)** | 0 |  | 11 (5.6%) | **8 (4.7%)** | 0 | 0 |
|  | **Ncrop-crop**<br>**hybrids (and %)** | 0 | **43**<br>**(89.5%**<br>**)** | **4 (1%)** | **Ncrop-**<br>**to-wild**<br>**hybrids**<br>**(and %)** | **15 (7.6%)** | **26 (18.1%)** | **0** | **0** |
| Hyrcanian (pink) | mean<br>introgression rate | 0 | **0.05** | 0 |  | 0 | **0.41** | 0 | 0 |

|  |  |  |  |  |  |  |  |  |  |  |
| --- | --- | --- | --- | --- | --- | --- | --- | --- | --- | --- |
|  | N (%) | 0 | **6**<br>**(12.5%)** | 0 |  | 0 | **34 (20.3%)** | 0 | 0 |  |
| Lorestan (light green) | mean introgression rate | 0 | 0.01 | 0 |  | 0 | 0.04 | 0 | 0 |  |
|  | N (%) | 0 | 2 (4%) | 0 |  | 0 | 6 (3.6%) | 0 | 0 |  |
| Wild Kurdistan (red) | mean introgression rate | 0 | **0.17** | 0 |  | 0 | **0.27** | 0 | 0 |  |
|  | N (%) | 0 | **12**<br>**(35%)** | 0 |  | 0 | **30 (17.9%)** | 0 | 0 |  |
| Western (orange) | mean introgression rate | 0.65 | 0.08 | 0 |  | **0.28** | 0.02 | 0.49 | 0.84 |  |
|  | N (%) | 1 (33%) | 3<br>(6.2%) | 0 |  | **33**<br>**(16.8%)** | 4 (2.3%) | 1<br>(20%) | 2<br>(33%) |  |
| Southern (brown) | mean introgression rate | 0 | 0.02 | 0 |  | **0.17** | 0.02 | 0.35 | 0 |  |
|  | N (%) | 0 | 2<br>(4.1%) | 0 |  | **13 (6.6%)** | 2 (1.1%) | 1<br>(20%) | 0 |  |
| Central (blue) | mean introgression rate | 0.32 | 0.02 | 0 |  | **0.46** | 0.01 | 0 | 0.11 |  |
|  | N (%) | 1 (33%) | 1 (2%) | 0 |  | **46**<br>**(23.4%)** | 1 (0.5%) | 0 | 1<br>(16.6%) |  |
|  | Nwild-to-crop hybrids (and %) | 2 (66%) | **26**<br>**(54.1%)** | 0 | Nwild-to-wild hybrids (and %) | **92**<br>**(46.9%)** | 77 (46.1%) | 2<br>(40%) | 3<br>(50%) |  |
|  | Total number of hybrids (and %) | **1** | **32**<br>**(66%)** | **2 (5%)** |  | **58 (28%)** | **54 (32%)** | **1**<br>**(20%)** | **2**<br>**(33%)** | **150**<br>**(32%)** |

We removed the 150 hybrids and all apple cultivars (Tables 2 and S2) and focused on gene flow within the “pure” Caucasian crabapple *M. orientalis*. We detected a significant but weak isolation by distance pattern across the Caucasus (*P* < 0.001, *R^2^* = 0.07, Figure S6), suggesting a high level of gene flow among the sampled geographic sites. We estimated *Sp* values for populations with at least five sampling sites and 20 individuals, *i.e.*, the Hyrcanian (pink) and the Central Armenian (blue) *M. orientalis* populations. *Sp* values were low but significant (*Sp* _Hyrcanian_pink_ = 0.0076, *Sp* _Central_blue_ == 0.0027, *P* < 0.001) suggesting a high level of historical gene flow within populations. However, the *Sp* value was higher for the Iranian population than for the Armenian population, suggesting a lower level of historical gene flow within the Hyrcanian (pink) population than the Central Armenian (blue) population. Contemporary estimates of gene flow further confirmed the occurrence of wild-to-wild, crop-wild and crop-crop gene flow in the Caucasus (Table S8 and Figure S7). Our results therefore are in agreement with the observations obtained above for the full dataset, suggesting substantial crop-crop, crop-wild and wild-wild gene flow in apples in Iran and the Caucasus.

### Additional domestication events from *M. orientalis* in Iran inferred with ABC-RF

We defined the populations used in the ABC framework from the clusters detected with STRUCTURE at *K*=12 for 550 wild and cultivated apple accessions, excluding admixed individuals (*i.e.*, with a membership coefficient < 0.85 to any given cluster, Figure S2, as recent gene flow can easily be seen from visual inspection of the bar plots). We excluded *M. baccata* as it was the most genetically differentiated species, and the Armenian cultivars as they were represented by only four individuals. We included two *M. sieversii* populations (from Kazakhstan and Kyrgyzstan), the cultivated (green and purple) populations from Iran, the standard *M. domestica* population and the seven populations of *M. orientalis* (four populations from Iran and three populations from Armenia, Table 1). We also simulated an ancestral population. In total, we therefore simulated thirteen populations (Figures S8 and S9).

To keep the number of scenarios tractable, we fixed the divergence histories of the wild apple populations as inferred from the NJ tree (Figures 2b, S8 and S9), assumed bidirectional gene flow among populations which is congruent with the high levels of crop-wild and crop-crop gene flow shown above. We built eight groups of scenarios to test whether the two Iranian cultivated apple populations diverged from the same wild population or whether they diverged from different populations. In order to assess the origin of the wild purple population, these eight groups of scenarios were simulated under two alternative evolutionary scenarios (Figures S8 and S9, respectively): the wild purple population was either assumed to have diverged from the ancestral population (Figure S8), or the Iranian wild and crop purple populations were assumed to be sister groups (Figure S9). A total of 46 scenarios was therefore simulated, representing 16 groups of hypotheses (see details in Figures S8 and S9).

For each step of the ABC-RF approach, the projection of the reference table and the observed datasets onto the two LDA axes that explained most of the variance of the summary statistics showed that the observed data fell within the distribution of the simulated summary statistics (Figure S10), forming distinct clouds for each scenario or groups of scenarios. Visual inspection of the LDA plots indicated that we had enough power to discriminate and select scenarios; results were subsequently validated by the ABC-RF inferences presented below.

We used a five-step nested ABC-RF approach. In the first step, we tested the divergence history of the purple wild and cultivated populations. All ten replicate ABC-RF analyses supported a sister relationship between the cultivated and wild purple populations (groups of scenarios 9 to 16, Figure S8, an average of 282 votes out of the 500 RF-trees; posterior probabilities = 65%, prior error rate = 0.98%, Table S9). In a second step, we compared the remaining groups (groups 9 to 16) to reconstruct the history of the Iranian cultivated apple populations. All ten replicate ABC-RF analyses supported that the two Iranian cultivated apple populations (green and purple) diverged from the same wild apple population (groups 9 to 12, an average of 296 votes out of the 500 RF-trees; posterior probabilities = 82%, prior error rate = 1.1%, Table S10). In the third step, we compared the remaining groups (groups 9 to 12) to test whether these Iranian cultivated apple populations diverged from *M. sieversii*, *M. domestica* or *M. orientalis*. All ten replicate ABC-RF analyses supported that the two Iranian cultivated apple populations originated from *M. orientalis* (groups 11 and 12, an average of 290 votes out of the 500 RF-trees; posterior probabilities = 79%, prior error rate = 0.2%, Table S11). In the fourth step, we compared the remaining groups (groups 11 and 12) to test whether the two Iranian cultivated apple populations were derived from *M. orientalis* from Iran or from Armenia. All ten replicate ABC-RF analyses supported that the two Iranian cultivated apple populations originated from *M. orientalis* from Iran (group 11, an average of 316 votes out of the 500 RF-trees; posterior probabilities = 92%, prior error rate = 0.10%, Table S12). For step five, we compared the last three remaining individual scenarios to select the final model. All ten replicate ABC-RF analyses supported that the two Iranian cultivated apple populations originated from the Iranian red *M. orientalis* population (sc45, an average of 316 votes out of the 500 RF-trees; posterior probabilities = 92%, prior error rate = 0.10%, Table S13). This nested ABC approach avoids comparing too many complex models with too many parameters, and is more powerful than testing all scenarios individually to disentangle the main evolutionary events characterizing demography and divergence (Estoup et al., 2018).

ABC-RF inferences therefore provided support for additional apple domestication events in Iran from the red *M. orientalis* population. The time of domestication was inferred to be *ca.* 826 years ago (confidence interval (CI): 130-5,030) for the green Iranian crop population and *ca*. 803 ya (CI: 160-3,969) for the purple crop population, respectively; by contrast *M. domestica* was inferred to have diverged from *M. sieversii ca*. 3,898 ya (CI: 270-7,670) (Table S14, Figure 3). The divergence time between the wild and the crop purple populations was estimated to be 418 ya (CI: 100-2,500). Note that the confidence intervals were relatively high for some parameters (*e.g.*, migration rates) as illustrated by the high normalized mean absolute error values (NMAE). High posterior probabilities and low *prior* error rates for model choice nevertheless indicate high support for the final model, even if some parameters such as migration rates cannot be precisely estimated.

**Figure 3.**
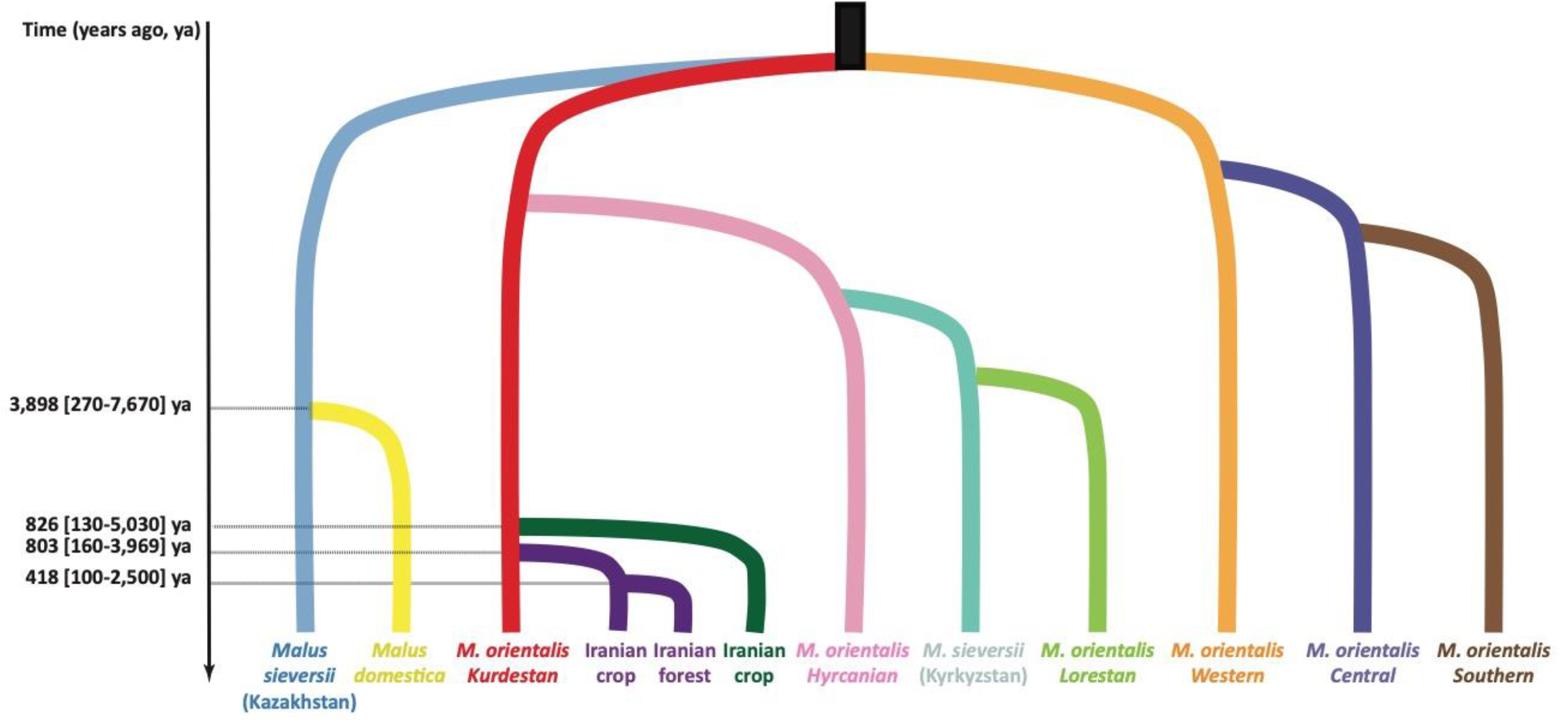
Most likely scenario of domestication of cultivated apples in Iran using random-forest approximate Bayesian computation (ABC-RF) combined with coalescent-based simulations (SC45, Figure S9). Population names correspond to the ones detected with STRUCTURE for *K* = 12, excluding admixed individuals (*i.e.*, individuals with a membership coefficient < 0.85 to any given cluster), *M. baccata* was excluded from the inferences. Bidirectional gene flow between populations was assumed. For clarity, only estimates of divergence time of the cultivated populations are provided, other parameters estimates are provided in Table S14.

### Range expansion and contraction of the wild apple *M. orientalis* in the Caucasus associated with the past glaciations

We investigated the spatial variation of genetic diversity and used ecological niche modeling to test the existence of past contraction and expansion of the range of the wild apple in the Caucasus. After removing the 150 crop-wild hybrids identified from the second STRUCTURE analysis, we found a significant positive correlation between longitude and allelic richness (Figure S11, average adjusted *R^2^* = 0.66, *P* < 0.0001) and a significant negative correlation between latitude and allelic richness (Figure S10, average adjusted *R^2^* = −0.43, *P* < 0.001). We also found that the Western (orange) population had the highest level of allelic richness (Tables 1 and S7 and Figure 4). The Western Caucasus may therefore have been a glacial refugium in the past. In addition to high levels of genetic diversity in the west, across northeastern Turkey and the Lesser Caucasus mountains in Armenia, we observed local hotspots of genetic diversity in the Hyrcanian Forests and the High Caucasus mountains (Figure 4), suggesting that these mountainous regions may have been potential glacial refugia.

**Figure 4.**
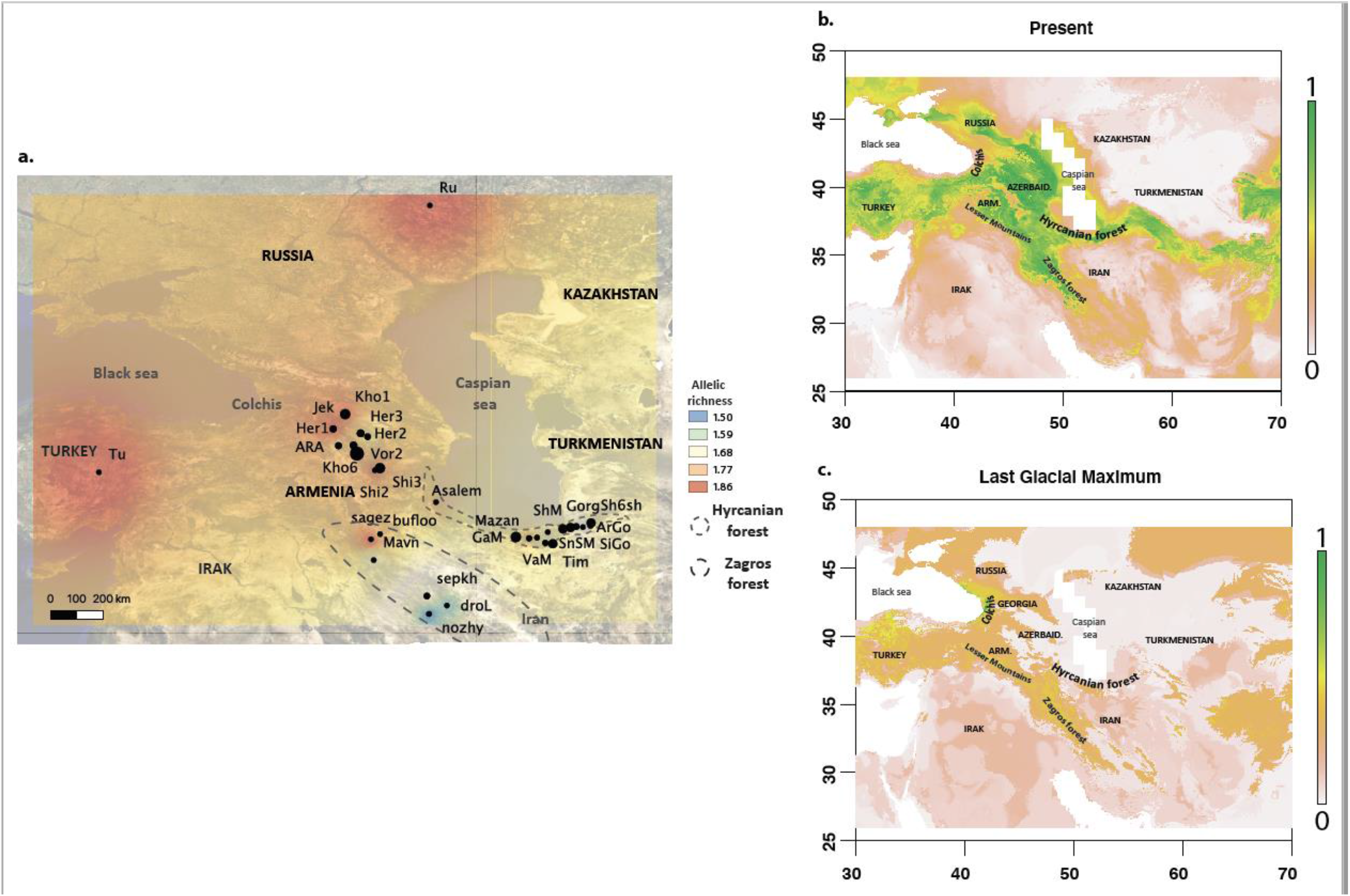
Spatial diversity and past contraction and expansion of *Malus orientalis* across the Caucasus. **a.** Spatial genetic diversity (allelic richness) at 36 sites (*N* = 339). **b. and c.** Ensemble forecasting of the three different algorithms (ANN, GLM and GAM) predicting the current and last glacial maximum (LGM) distribution range of suitable areas for *M. orientalis*, respectively. The probabilities of being a suitable habitat are given in the legend. The Colchis and Hyrcanian regions are shown on the maps.

Ecological niche modeling further indicated past contraction and expansion of the *M. orientalis* range. Model performance as assessed with AUC and TSS was high (Table S15), indicating that the ANN, GLM and GAM algorithms fitted the data well (Allouche, Tsoar, & Kadmon, 2006; Fieldings & Bell, 1997; Monserud & Leemans, 1992). The following six bioclimatic variables were found to have high predictive power: mean diurnal range temperature (bio2), temperature seasonality (bio4), minimum temperature of the wettest quarter (bio8), minimum temperature of the driest quarter (bio9), annual precipitation (bio12) and precipitation of the coldest quarter (bio19). These bioclimatic variables were used to calibrate the models to predict the past and present distribution of *M. orientalis*. The MIROC model (Figure 4) predicted that the areas suitable for *M. orientalis* during the LGM contracted to the western Lesser Caucasus and northeastern Turkey along the Black Sea and into the Colchis region, and also in the eastern part of the Hyrcanian Forests, near Azerbaijan, in agreement with the genetic data (Figure 4). The climatic model therefore suggested that populations of the Caucasian wild apple *M. orientalis* may have been maintained in at least two glacial refugia.

## Discussion

Our study provides insights into the natural and anthropogenic divergence history of apples in a hotspot of crop diversity, the Caucasus. Our results revealed that the evolution of the domesticated apple involved an additional wild species, *M. orientalis*. We identified two distinct cultivated gene pools in Iran that were well differentiated from the standard *M. domestica* apple cultivars and were derived from the same wild Iranian *M. orientalis* population. In addition, several cultivated apple genotypes from the Caucasus were found to belong to the Caucasian *M. orientalis* gene pool, suggesting that local farmers use the Caucasian crabapple for cultivation, which may represent the early stages of domestication. *Malus orientalis* has also contributed to the Caucasian cultivated apple germplasm through wild-to-crop introgression; a similar process has been previously described in apples in Europe (Cornille et al., 2014, 2012). Conversely, crop-to-wild gene flow, which has been reported in Europe, was also detected in the Caucasus (Cornille et al., 2015). We also showed that « pure » *M. orientalis* in this region displays a clear spatial genetic structure with at least seven populations spread across the Caucasus. The combination of niche modeling and population genetics approaches suggested that these populations resulted from range contraction and expansion associated with the past glaciations. Iran is therefore an additional center of apple domestication, where distinct cultivated gene pools evolved from their progenitor *M. orientalis*. *Malus orientalis* has also contributed to cultivated gene pools in the Caucasus and Iran through wild-to-crop gene flow. Our results thereby reinforce the view that apple tree domestication has involved multiple wild apple species from multiple geographic regions, a pattern that has also been found in apricot and pear trees (Groppi, Liu, Cornille, Decroocq, & Decroocq, 2021; Liu et al., 2019; Wu et al., 2018). We also found evidence of substantial hybridization between domestic and wild forms, and this has also been described in other fruit trees (Groppi et al., 2021; Liu et al., 2019; Wu et al., 2018). In addition, this study highlights the impact of climate change on the natural divergence of a wild fruit tree and provides a starting point for apple conservation and breeding programs in the Caucasus and Iran.

### Additional events of apple domestication from *M. orientalis* in Iran

The occurrence of two distinct cultivated populations in Iran, which are genetically differentiated from the cultivated apple *M. domestica* and are derived from *M. orientalis*, suggests that Iran is an additional center of apple domestication. Domesticated populations are expected to be nested within their source population because they diverged recently from a subset of individuals within the source population (Matsuoka et al., 2002). However, the green and purple Iranian cultivated populations were not nested within any wild population (unlike *M. domestica* that was nested within *M. sieversii*). Nevertheless, the monophyly of the two Iranian cultivated groups (green and purple) suggests that they diverged from the same wild population, perhaps corresponding to two successive domestication steps, or represent independent domestication events from the same progenitor. We tested these hypotheses and inferred more precisely the Iranian cultivated apple domestication history using coalescent-based methods combined with approximate Bayesian computation. This confirmed that the two Iranian cultivated populations diverged from a *M. orientalis* population in Iran, most likely from the Kurdestan province (red population). Thus, despite the spread of the cultivated apple *M. domestica* along the Silk Trade Routes that crossed Iran and the South Caucasus to reach Turkey (Canepa, 2010; Spengler, 2019), specific domestication events in this region have resulted in local cultivated apple gene pools. It is still not clear, however, whether these two cultivated apple populations represent *de novo* domestication events. Distinct genetic ancestries may not reflect independent *de novo* domestication, but may instead represent a single domestication event with multiple origins (Choi et al., 2017; Gros-Balthazard & Flowers, 2021). The occurrence of independent domestication events in many crop species is a source of ongoing debate (Besnard et al., 2018; Choi et al., 2017; Gros-Balthazard & Flowers, 2021). In apricot and pear trees, there is evidence of independent domestication events in Europe and Asia, with distinct cultivated gene pools of different ancestries which display different genomic regions showing footprints of selection (Groppi et al., 2021; Liu et al., 2019; Wu et al., 2018). Here, our data shows that there are distinct cultivated apple populations, with different geographic origins and which are derived from distinct ancestral gene pools. Studies of the regions under selection during domestication in the Iranian and European cultivated apple genomes will likely provide insights into whether multiple *de novo* domestication events occurred. Independent selection regimes in each of the cultivated genetic groups would be a hallmark of multiple domestications, as shown in apricot and pear trees (Groppi et al., 2021; Wu et al., 2018). In addition, coalescent-based methods combined with ABC-RF provided a powerful statistical framework for inferring the apple domestication history in the Caucasus and Iran. However, a broader view of the domestication history of the apple tree will require additional sampling of *M. sylvestris*, which is another contributor to the *M. domestica* gene pool (Cornille et al., 2014, 2012), as well as of local varieties across Eurasia, which are missing in this study.

### Wild-to-crop gene flow from *M. orientalis* to local cultivars and cultivation of the local wild apple in the Caucasus

The Caucasian crab apple has considerably contributed to the Caucasian apple germplasm through wild-to-crop introgression. We found evidence of substantial wild-crop and crop-crop gene flow in the Caucasus. Indeed, we found that 41.6% of Iranian cultivars were introgressed by local wild apple gene pools or were an admixture of two cultivated gene pools. This extensive wild-to-crop and crop-crop gene flow is strikingly similar to the pattern documented in apples in Europe. *Malus sylvestris* has been shown to be a significant contributor to the *M. domestica* gene pool through recurrent and recent hybridization and introgression events ever since the cultivated apple was introduced in Europe by the Greeks around 1,500 years ago (Cornille et al., 2012). Conversely, substantial crop-to-wild gene flow has been reported from *M. domestica* to *M. sylvestris* (Cornille et al., 2015). Similarly, we found many crop-to-wild hybrids (*M. orientalis* introgressed with *M. domestica*) in the forests of Armenia and Iran. Extensive gene flow has been found during the domestication of other fruit trees (Arroyo-García et al., 2006; Cornille et al., 2012; Decroocq et al., 2016; Diez et al., 2015; Duan et al., 2017; Liu et al., 2019; Meyer, Duval, et al., 2012; Myles et al., 2011). The evolutionary consequences of crop-to-wild gene flow remains unclear in fruit trees (Feurtey, Cornille, Shykoff, Snirc, & Giraud, 2017a); the extent to which crop-to-wild gene flow may threaten the Caucasian crab apple remains to be tested.

Our findings also suggest that farmers in the Caucasus and Iran grow *M. orientalis* rather than *M. domestica*. Indeed, the four Armenian cultivars share their gene pools with the Western (orange) and Central (blue) Caucasian crabapple populations, and some Iranian cultivars are fully assigned to wild populations (red and purple) in Iran. Coalescent based methods suggest that the wild purple population found in the forest is feral or results from a recent domestication event. It is difficult to discriminate between these two hypotheses using only genotypes as both predict a sister relationship between the wild and cultivated purple populations. Individual isolated trees belonging to the purple population are found in natural mountainous areas close to fruit trees grafted on *Crataegus*, although they are not themselves cultivated (personal observation. H. Yousefzadeh). In addition, the wild purple population had a lower level of genetic diversity than the other wild Iranian populations. Altogether, this may suggest that this population has recently escaped from cultivation. Like domestication, feralization can be seen as a process that is accompanied by admixture and introgression, and can be accompanied by a range of genetic, phenotypic and demographic changes (Mabry, Rowan, Pires, & Decker, 2021). Feral populations have also been documented in other fruit trees, including olive (Besnard et al., 2018), almond (Balaguer-Romano et al., 2021) and apricot (Spengler, Chang, & Tourtellotte, 2013). However, the trees found in the Hyrcanian Forests could also be the result of a first domestication step, as wild apples are widely used for cultivation in Iran (personal communication H. Yousefzadeh and from our results). Additional phenotypic data to identify shifts in phenotypic traits associated with domestication or feralization are required to further disentangle these hypotheses. The cultivation of wild genotypes (here, from the red or purple populations) would not be surprising as *M. orientalis* can grow in mountainous areas, is highly resistant to pests, diseases and drought (Amirchakhmaghi et al., 2018; Büttner, 2001; Höfer et al., 2013; Volk et al., 2008), and has high-quality fruits that have several features that are intermediate between those of *M. sylvestris* and *M. sieversii* (Cornille et al., 2014). The use of local wild apples has also been documented in Europe for specific purposes at different times in history (Tardío, Arnal, & Lázaro, 2020).

### The genetic relationships between the two wild contributors are still unclear

Understanding of the genetic relationships among wild populations that are closely related to a crop gene pool is crucial to better understand crop domestication. Here, we found that *M. sieversii* from Kazakhstan was a sister group to the *M. domestica* gene pool, while *M. sieversii* from Kyrgyzstan was closer to *M. orientalis* from the Zagros and Hyrcanian regions. *Malus sieversii* and *M. orientalis* are referred as to different species, but our results suggest that *M. orientalis* may be polyphyletic and intermingled with *M. sieversii* (Harris et al., 2002; Nikiforova, Cavalieri, Velasco, & Goremykin, 2013; Robinson, Harris, & Juniper, 2001). The relationship between *M. sieversii* and *M. orientalis* is therefore still unclear. Deciphering the relationship between *M. sieversii* and *M. orientalis*, and even the species status of *M. orientalis,* is not only a taxonomic exercise but is needed to better understand apple domestication. This question needs to be resolved urgently, *M. sieversii* being endangered across its distribution (Omasheva et al., 2017; Zhang, Li, & Li, 2018), as confirmed here by the lower level of genetic diversity of the *M. sieversii* population in Kyrgyzstan.

### The natural divergence history of the Caucasian wild apple was shaped by the past glaciations

The climatic variations since the last glacial maximum, along with the landscape features of the Caucasus, have likely shaped the population structure and diversity of the Caucasian wild apple. We identified seven populations of *M. orientalis* in the Caucasus and Iran: one highly genetically differentiated population in the Western Caucasus (Turkey, Russia and northwestern Armenia), two in Armenia (a southern and a Central population) and four in Iran, including two in the Zagros Forest (one in the Kurdestan province and one in the Lorestan province) and two in the Hyrcanian Forests bordering the southern Caspian Sea. These wild apple populations likely arose from isolation in several refugia during the last glacial maximum. This hypothesis is supported by the observation of a large hotspot of genetic diversity located in the Western Caucasus, and several local hotpots of genetic diversity in Armenia and the Hyrcanian Forests (Zazanashvili et al., 2020). Ecological niche modeling further supported the existence of strong contractions in the range of *M. orientalis* in the Western Caucasus bordering the Black Sea (including the Colchis region), as well as in the Lesser Caucasus and in some parts of the Hyrcanian Forests. Additional samples from the Western Caucasus are required to confirm this hypothesis. These glacial refugia have been described in relation to other species (Parvizi et al., 2019). Indeed, two refugia are recognized in the Caucasus (Tarkhnishvili *et al.* 2012; Yousefzadeh *et al.* 2012; Bina *et al.* 2016; Aradhya *et al.* 2017): a major forest refugium between the western Lesser Caucasus and northeastern Turkey (including the Colchis region in the catchment basin of the Black Sea) and the Hyrcanian refugium at the southern edge of the Caucasus. Further sampling of *M. orientalis* in the far Western and Eastern Caucasus and genotyping with the same microsatellite markers are needed to uncover the role of these two refugia in the divergence history of *M. orientalis*.

We also found that the natural divergence history of the Caucasian wild apple involved gene flow across the Caucasus. The weak but significant isolation-by-distance pattern further supports the existence of substantial gene flow among wild apple populations in the Caucasus. Widespread gene flow during divergence associated with the last glacial maximum has been documented for another wild apple relative *M. sylvestris* (Cornille et al., 2013). Calculation of the *Sp* parameter within the largest populations revealed high levels of historical gene flow within populations. *Sp* can also be used to compare the dispersal capacities of *M. orientalis* with that of other plants (Cornille et al., 2013; Cornille, Gladieux, & Giraud, 2013; Vekemans & Hardy, 2004). The Caucasian wild apple showed dispersal capacities that were similar to previous estimates in other wild apple species and lower than that of wind-dispersed trees. Wild apples can disperse over kilometers (Cornille et al., 2015; Feurtey, Cornille, Shykoff, Snirc, & Giraud, 2017). The spatial population structure was somewhat stronger in Iran than in Armenia, suggesting lower levels of gene flow in the Hyrcanian population. In addition to having a stronger genetic structure, the Iranian populations had lower genetic diversity then the Armenian populations, especially the Zagros and Kurdestan populations. In Iran, traditional animal husbandry is a widespread practice (Soofi et al., 2018). Such intensive farming environments may lead to forest fragmentation and may impact wild apple populations, which form low density populations. The future of Iranian wild apple populations, especially in the south where genetic diversity is low, may depend on our ability to protect them through conservation programs.

## Conclusion

We identified Iran as a key center in the evolution and domestication of apple, and *M. orientalis* as an additional contributor to the evolutionary history of cultivated apples in Iran and in the Caucasus. We also provided insights into the processes underlying the natural divergence of this emblematic wild species and identified several populations that could be the target of conservation programs. However, *M. orientalis* in the Caucasian ecoregion is highly diverse and further investigations and additional sampling are necessary, as well as a better assessment of its species status and genetic relationship with *M. sieversii*. Indeed, a better understanding of the properties of functional genetic diversity and of the ecological relationships of wild apples in their ecosystem are needed for developing and implementing effective conservation genetic strategies in this region (Teixeira & Huber, 2021). Our study revealed the role of gene flow and human practices in natural and anthropogenic divergence processes of an emblematic fruit tree in the Caucasus and Iran. Our results are consistent with those reported for other woody perennials, including apricot (Groppi et al., 2021; Liu et al., 2019), olive (Besnard et al., 2018; Diez et al., 2015), pear (Volk & Cornille, 2019; Wu et al., 2018), and date palm (Flowers et al., 2019). This study also supports the view that domestication of fruit trees was likely a geographically diffuse and protracted process, involving multiple, geographically disparate origins of domestication (Groppi et al., 2021; Wu et al., 2018).

## Acknowledgements

We thank the Franco-Iranian Campus France program « Gundhishapur » 2016-2018, the Institut Diversité Écologie et Évolution du Vivant (IDEEV) and ATIP-Avenir for funding. We thank Bolotbek Tagaev (Sustainable Livelihoods Coordinator of FFI-Kyrgyzstan) for sampling and prospection, Fauna & Flora International and more specifically the Global Trees Campaign (GTC) Program. We also thank Adrien Falce, Olivier Langella and Benoit Johannet for help and support on the INRAE-Génétique Quantitative et Evolution-Le Moulon lab cluster and the genotyping platform GENTYANE INRA UMR 1095. We thank the INRAE MIGALE bioinformatics platform (http://migale.jouy.inra.fr) for providing help and support, in particular Véronique Martin, Eric Montaubon and Valentin Loux. We also thanks Céline Bellard for her advices for ecological niche modeling analyses.

## Data Availability

SSR data are available on the DRYAD repository XXXX.

## Author Contributions

AC, HY conceived and designed the experiments; AC, HY obtained funding; HB, HY, SF, HBa, IG, AN, AC, JS, DG, AK sampled the material; AV, CR, AR, MF performed the molecular work; AC, HB analyzed the data; AC, HB, HY: wrote the original draft and preparation of the figures; AC, HB, HY, TG, XC, IG, AN and all co-authors: gave critical inputs in final draft and revisions.

