## Supporting information for "Evidence of an additional wild contributor, *Malus orientalis* Uglitzk., to the genome of cultivated apple varieties in the Caucasus and Iran"

Article acceptance date: NA

The following Supporting Information is available for this article:

**Table S1.** Passport information and genotypes for the 26 microsatellite markers of the wild and cultivated apples used in this study (*N*=550), with their geographic or germplasm origins, when applicable, provider(s), geographic coordinates (X and Y), membership coefficients inferred with STRUCTURE for *K*=12, population assignation (*i.e.*, membership > 0.85 to a given cluster), and when available width and length fruit size for the Iranian cultivars.

**Table S2.** Membership coefficient inferred with STRUCTURE (excluding *M. sieversii* and *M. baccata, N*=466) for *K*=9 used to detect crop-wild hybrids in the Caucasus.

**Table S3**. Prior distributions for each model parameter used for Random Forest approximate Bayesian computations to reconstruct the domestication history of the Iranian apple.

**Table S4.** Presence data point (57) of *Malus orientalis* detected as pure (*i.e.,* individual assigned with a membership coefficient >0.9 to a wild apple gene pool in the second STRUCTURE analysis excluding *M. baccata* and *M. sieversii*) used for ecological niche modeling in the Caucasus.

**Table S5.** Correlation among the 19 bioclimatic variables used for the ecological niche modelling for *Malus orientalis* in the Caucasus.

**Fig. S1.** Delta *K,* and probability for each *K*, plotted against *K* values inferred with STRUCTURE for the two datasets: all samples (*N*=550, a and b), and excluding *Malus baccata* and *Malus sieversii* (*N*=466, c and d).

**Fig. S2.** Bayesian clustering obtained with STRUCTURE for the wild and cultivated apple using 26 microsatellite markers, from *K*=2 to *K*=13, based on the whole dataset (*N*=550).

**Fig. S3.** Distribution of the maximum membership coefficients inferred with STRUCTURE for the wild and cultivated apples (*N*=550, 26 SSR markers) at *K*=12. The vertical line at 0.85 represents the threshold used to assign an individual to a given cluster. Individuals with a membership coefficient > 0.85 were considered as admixed genotypes (in grey in other figures).

**Table S6.** Pairwise genetic differentiation estimates (*F_ST_*, lower triangle, and Jost’s *D,* upper triangle) between the twelve populations (*i.e.*, groups including individuals assigned with a membership coefficient > 0.85 to a given cluster) detected with STRUCTURE at *K*=12 (*N* = 424, 26 microsatellite markers). All estimates were significant, except between the wild and cultivated purple populations.

**Table S7.** Adjusted *p-values* from the Wilcoxon signed rank test to compare allelic richness (*A_R_*, lower triangle) and private allelic diversity (*A_P_*, upper triangle) among the twelve wild and cultivated apple populations detected with the second STRUCTURE analysis at *K*=12. Only comparisons in bolded were significant (*P* < 0.05).

**Figure S4.** Bayesian clustering obtained with STRUCTURE for the wild and cultivated apple using 26 microsatellite markers, from *K*=2 to *K*=10, based on the dataset excluding *M. sieversii* and *M. baccata* (*N*=466).

**Figure S5.** Distribution of the maximum membership coefficient inferred with STRUCTURE for the 446 wild and cultivated apples at *K*=9.

**Figure S6.** Pairwise genetic differentiation (*F_ST_/(1- F_ST_)*) in function of the logarithm of the distance for *Malus orientalis* (*N*=339, 36 sample sites with at least four individuals) in the Caucasus.

**Table S8.** **Matrix of gene flow inferred with BayesAss** among apple populations in the Caucasus and Iran.

**Figure S7**. Circle plot representing gene flow among populations in the Caucasus and Iran inferred with BayesAss.

**Figure S8**. Details of the scenarios tested with approximate Bayesian computation for reconstructing the domestication history of the Iranian cultivated apple*.*

**Figure S9**. Details of the scenarios tested with approximate Bayesian computation for reconstructing the domestication history of the Iranian cultivated apple*.*

### **Figure. S10.** Linear discriminant analysis, and associated variance plot, obtained for each of the five random forest approximate Bayesian computation steps to infer the Iranian cultivated evolutionary history.

**Table S9.** Results of the ABC-RF algorithm inferring the origin of the wild Iranian apple population (ABC-RF step 1).

**Table S10.** Results of the ABC-RF algorithm inferring whether the two cultivated Iranian populations are a sister group which diverged from a same wild population (groups 9 to 12) or diverged from two different wild populations (groups 13 to 16, Figure S5) (ABC-RF step 2).

**Table S11.** Results of the ABC-RF algorithm inferring whether sister cultivated apple populations originated from *M. sieversii/M.domestica* (groups 9 and 10) or *M. orientalis* (groups 11 and 12) (ABC-RF step 3).

**Table S12.** Results of the ABC-RF algorithm infering whether sister cultivated apple populations originated from *M. orientalis* from Iran (group 11) or from Armenia (group 12) (ABC-RF step 4).

**Table S13.** Results of the ABC-RF algorithm inferring whether the sister Iranian cultivated apple populations originated from one of the three Iranian *M. orientalis* populations (ABC-RF step 5).

**Table S14.** Parameter estimates infered for the Iranian cultivated apple domestication history, computed with the most likely (SC45, Figure S5).

**Fig. S11.** Spatial genetic variation for *Malus orientalis* (*N*=339, 36 sample sites) in the Caucasus.

**Table S15.** AUC (ROC curve) and TSS indices for bioclimatic variables for the modeling algorithms (ANN, GAM, GLM) and repetitions for *Malus orientalis.*

**Table S1.** Passport information and genotypes for the 26 microsatellite markers of the wild and cultivated apples used in this study (*N*=550), with their geographic or germplasm origins, when applicable, provider(s), geographic coordinates (X and Y), membership coefficients inferred with STRUCTURE for *K*=12, population assignation (*i.e.*, membership > 0.85 to a given cluster), and when available width and length fruit size for the Iranian cultivars.

See excel file Table S1.xlsx

**Table S2.** Membership coefficient inferred with STRUCTURE (excluding *M. sieversii* and *M. baccata, N*=466) for *K*=9 used to detect crop-wild hybrids.

See excel file Table S2

**Table S3.** Prior distributions for each model parameter used for Random Forest approximate Bayesian computations to reconstruct the domestication history of the Iranian apple. Prior distributions are uniform or log-uniform between lower and upper bounds.

| **Parameter** | **Distribution** | **Lower bound** | **Upper bound** |
| --- | --- | --- | --- |
| *N_ANC_* | *unif* | 50 | 200 |
| *N_X_* | *unif* | 50 | 100 |
| *T_WILD_X_* (in years) | *logunif* | 30,000 | 300,000 |
| *T_CROP_X_* (in years) | *logunif* | 100 | 10,000 |
| *μ* | *logunif* | 10^-4^ | 10^-3^ |
| *m_x-y_* | *unif* | 10^-6^ | 10^-4^ |

*N_ANC_*: effective population size of the ancestral population*; N_X_*: effective population size of population *X*; *T_CROP_*: divergence time of the *X* crop population; *T_W_*: divergence time of the *W* wild population*;* *m_x-y_*: migration rate per generation between *x* and *y* populations, with *x* and *y* referred to as population identification number + 1 presented in Figures S9 and S10; *μ*: mutation rate; unif: uniform distribution; *log-unif*: log-uniform distribution.

**Table S4.** Presence data point (57) of *Malus orientalis* detected as pure (*i.e.*, individual assigned with a membership coefficient >0.9 to a wild apple gene pool in the second STRUCTURE analysis excluding *M. baccata* and *M. sieversii*) used for ecological niche modeling in the Caucasus.

| Longitude | Latitude |
| --- | --- |
| 46.0208333333333 | 41.4791666666667 |
| 44.6458333333333 | 41.3125 |
| 45.1875 | 41.3125 |
| 45.1875 | 41.2708333333333 |
| 44.7291666666667 | 40.7291666666667 |
| 44.7291666666667 | 40.6875 |
| 45.8125 | 40.5208333333333 |
| 45.7708333333333 | 40.4791666666667 |
| 46.1041666666667 | 40.3958333333333 |
| 44.9375 | 40.0208333333333 |
| 45.5208333333333 | 40.0208333333333 |
| 45.6875 | 39.7291666666667 |
| 45.6875 | 39.6875 |
| 46.5625 | 39.1041666666667 |
| 46.6041666666667 | 39.1041666666667 |
| 46.6458333333333 | 39.1041666666667 |
| 46.6875 | 39.1041666666667 |
| 46.3958333333333 | 39.0625 |
| 46.3958333333333 | 39.0208333333333 |
| 48.8541666666667 | 37.7708333333333 |
| 55.1458333333333 | 36.8958333333333 |
| 54.3958333333333 | 36.7708333333333 |
| 54.5208333333333 | 36.7708333333333 |
| 55.1041666666667 | 36.7708333333333 |
| 54.3125 | 36.7291666666667 |
| 54.6458333333333 | 36.6875 |
| 53.9791666666667 | 36.6458333333333 |
| 54.0625 | 36.6458333333333 |
| 53.3958333333333 | 36.5625 |
| 46.6041666666667 | 36.4791666666667 |
| 52.1041666666667 | 36.3541666666667 |
| 52.1458333333333 | 36.3541666666667 |
| 52.4375 | 36.3541666666667 |
| 52.0625 | 36.3125 |
| 52.9375 | 36.3125 |
| 46.2291666666667 | 36.2708333333333 |
| 52.6458333333333 | 36.2708333333333 |
| 53.2708333333333 | 36.1041666666667 |
| 53.6041666666667 | 36.0625 |
| 53.6458333333333 | 36.0625 |
| 46.2291666666667 | 35.4375 |
| 46.3541666666667 | 35.3958333333333 |
| 48.4375 | 33.9791666666667 |
| 48.4791666666667 | 33.9791666666667 |
| 48.4375 | 33.9375 |
| 48.4791666666667 | 33.9375 |
| 48.2708333333333 | 33.8958333333333 |
| 48.9375 | 33.8958333333333 |
| 48.1875 | 33.8541666666667 |
| 48.2708333333333 | 33.8541666666667 |
| 49.3125 | 33.5625 |
| 49.3541666666667 | 33.5625 |
| 48.7291666666667 | 33.3958333333333 |
| 48.4375 | 33.2291666666667 |
| 48.5625 | 33.2291666666667 |
| 48.6041666666667 | 33.2291666666667 |
| 49.3125 | 33.2291666666667 |

**Table S5. Correlation among the 19 bioclimatic variables used for the ecological niche modeling for *Malus orientalis* in the Caucasus.** Red and green colors indicate correlations > 0.75.

|  | bio_1 | bio_2 | bio_3 | bio_4 | bio_5 | bio_6 | bio_7 | bio_8 | bio_9 | bio_10 | bio_11 | bio_12 | bio_13 | bio_14 | bio_15 | bio_16 | bio_17 | bio_18 | bio_19 |
| --- | --- | --- | --- | --- | --- | --- | --- | --- | --- | --- | --- | --- | --- | --- | --- | --- | --- | --- | --- |
| bio_1 | 1.00 |  |  |  |  |  |  |  |  |  |  |  |  |  |  |  |  |  |  |
| bio_2 | 0.29 | 1.00 |  |  |  |  |  |  |  |  |  |  |  |  |  |  |  |  |  |
| bio_3 | 0.48 | 0.85 | 1.00 |  |  |  |  |  |  |  |  |  |  |  |  |  |  |  |  |
| bio_4 | 0.04 | 0.34 | -0.16 | 1.00 |  |  |  |  |  |  |  |  |  |  |  |  |  |  |  |
| bio_5 | 0.92 | 0.58 | 0.61 | 0.32 | 1.00 |  |  |  |  |  |  |  |  |  |  |  |  |  |  |
| bio_6 | 0.98 | 0.14 | 0.43 | -0.15 | 0.83 | 1.00 |  |  |  |  |  |  |  |  |  |  |  |  |  |
| bio_7 | -0.11 | 0.75 | 0.29 | 0.80 | 0.27 | -0.31 | 1.00 |  |  |  |  |  |  |  |  |  |  |  |  |
| bio_8 | 0.03 | -0.66 | -0.36 | -0.53 | -0.25 | 0.18 | -0.72 | 1.00 |  |  |  |  |  |  |  |  |  |  |  |
| bio_9 | 0.90 | 0.39 | 0.44 | 0.29 | 0.91 | 0.83 | 0.13 | -0.30 | 1.00 |  |  |  |  |  |  |  |  |  |  |
| bio_10 | 0.98 | 0.36 | 0.45 | 0.25 | 0.97 | 0.92 | 0.07 | -0.09 | 0.93 | 1.00 |  |  |  |  |  |  |  |  |  |
| bio_11 | 0.98 | 0.22 | 0.52 | -0.16 | 0.85 | 0.99 | -0.26 | 0.13 | 0.82 | 0.92 | 1.00 |  |  |  |  |  |  |  |  |
| bio_12 | -0.56 | -0.30 | -0.22 | -0.46 | -0.62 | -0.46 | -0.26 | 0.21 | -0.68 | -0.64 | -0.45 | 1.00 |  |  |  |  |  |  |  |
| bio_13 | -0.42 | -0.09 | 0.02 | -0.49 | -0.44 | -0.33 | -0.19 | 0.11 | -0.53 | -0.50 | -0.30 | 0.95 | 1.00 |  |  |  |  |  |  |
| bio_14 | -0.76 | -0.70 | -0.73 | -0.25 | -0.87 | -0.67 | -0.33 | 0.54 | -0.85 | -0.79 | -0.70 | 0.61 | 0.38 | 1.00 |  |  |  |  |  |
| bio_15 | 0.53 | 0.86 | 0.88 | 0.12 | 0.70 | 0.43 | 0.44 | -0.62 | 0.59 | 0.55 | 0.51 | -0.28 | -0.01 | -0.88 | 1.00 |  |  |  |  |
| bio_16 | -0.42 | -0.06 | 0.01 | -0.42 | -0.43 | -0.35 | -0.12 | 0.04 | -0.52 | -0.49 | -0.32 | 0.95 | 0.99 | 0.37 | 0.01 | 1.00 |  |  |  |
| bio_17 | -0.73 | -0.72 | -0.73 | -0.28 | -0.86 | -0.63 | -0.38 | 0.57 | -0.83 | -0.77 | -0.67 | 0.61 | 0.38 | 1.00 | -0.90 | 0.36 | 1.00 |  |  |
| bio_18 | -0.66 | -0.50 | -0.35 | -0.60 | -0.80 | -0.53 | -0.45 | 0.61 | -0.90 | -0.76 | -0.53 | 0.72 | 0.60 | 0.82 | -0.59 | 0.56 | 0.82 | 1.00 |  |
| bio_19 | 0.49 | 0.52 | 0.47 | 0.24 | 0.61 | 0.41 | 0.33 | -0.60 | 0.59 | 0.53 | 0.45 | 0.04 | 0.22 | -0.66 | 0.74 | 0.28 | -0.66 | -0.61 | 1.00 |

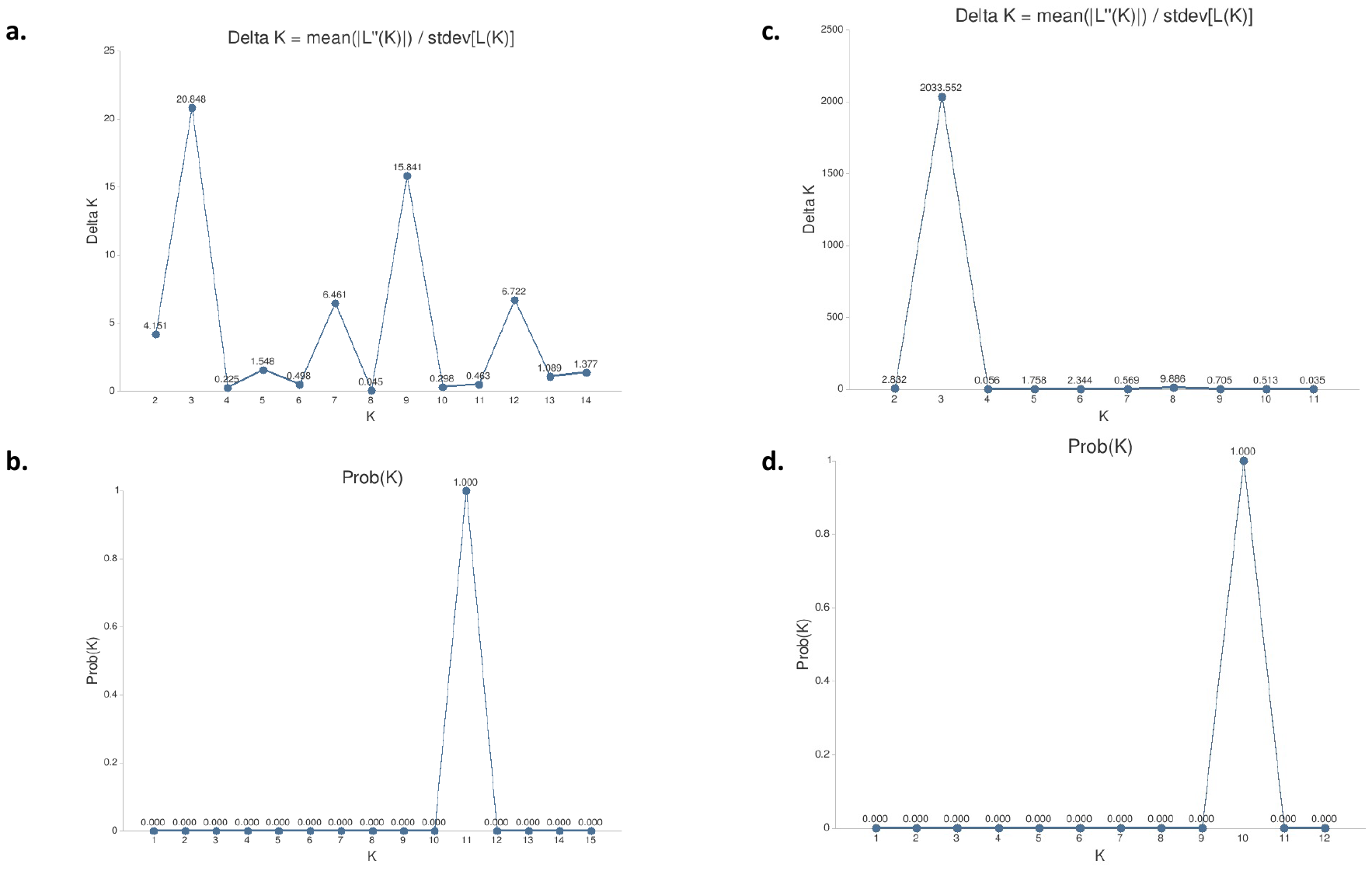
**Figure S1. Delta *K,* and probability for each *K*, plotted against *K* values inferred with STRUCTURE for the two datasets**: all samples (*N*=550, a and b), and excluding *Malus baccata* and *Malus sieversii* (*N*=466, c and d).

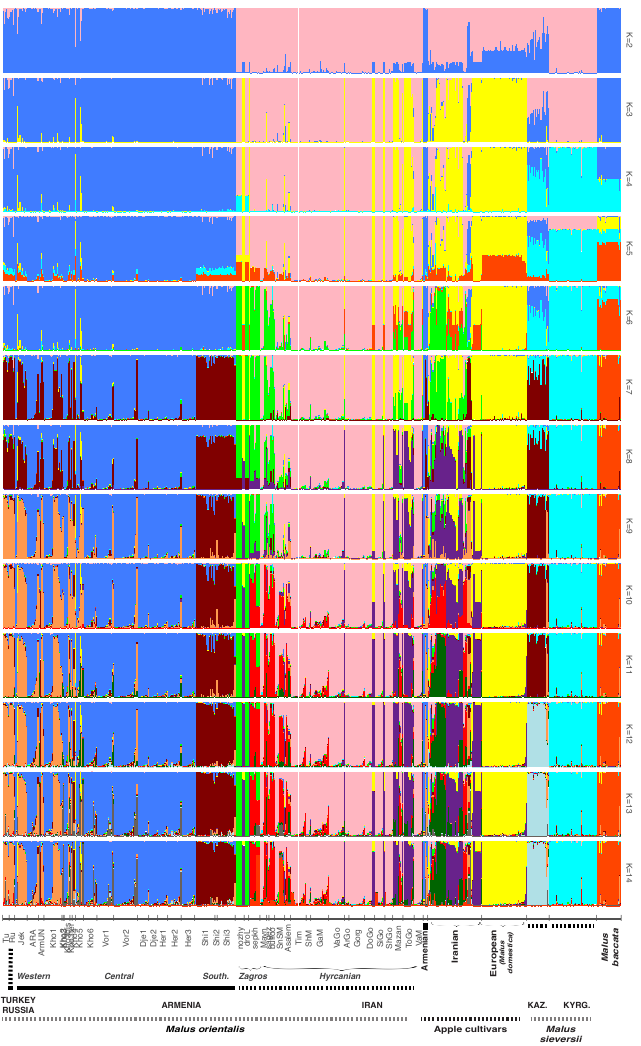

**Figure S2.** **Bayesian clustering obtained with STRUCTURE for the wild and cultivated apple using 26 microsatellite markers, from *K*=2 to *K*=14, based on the whole dataset (*N*=550).** Each individual is represented by a vertical bar, partitioned into *K* segments representing the proportions of ancestry of its genome in *K* clusters. Armenian sites (Shi1: Shikahogh1, Shi2: Shikahogh2, Shi3: Shikahogh3, Kho1: KhosrovReserve1, Kho2: KhosrovReserve2, Kho3: KhosrovReserve3, Kho3bis: KhosrovReserve3bis; Kho3ter: KhosrovReserve3ter, Kho4: KhosrovReserve4, Kho5: KhosrovReserve5, Kho6: KhosrovReserve6, Vor2: Vorotanpass2, Vor1: Vorotanpass1, Dje2: Djermuk2, Dje1: Djermuk1, Her2: Hermon2, Her1: Hermon1, Her3: Hermon3, ARA: ARA, Jek: Jermouck, Arm_unknown: Arm_unknown), Iranian sites (nozhy: Nozhyan region , DroL: Drood region, Sepkh: Sepedkoh region, Mavn: Marivan region, Sagez: Sagez region, Bufloo: Bufloo region, Asalem: Asalem region, Tim: Tim region; Tilak: Tilak region, SnSM: Sangdeh region, ShM: Shit region, VaM: Vaz region, GaM: Ganasara region, Mazan: Mazan region; Amol: Amol abesk region , VaGo: Vatna region, ArGo: Arsam region, Gorg: Gorgan province, DoGo: Dorak region, ToGo: Tokestan region, SiGo: Siamarzkoh region, ShGo: Sheshab region), DOM : *Malus domestica*, bacc: *Malus baccata*, siev: *Malus sieversii*. The main regions in the Caucasus and/or defined in the main text are also added: Western Caucasus (Turkey and Russia), Central and Southern Armenia, Zagros and the Hyrcanian forest.

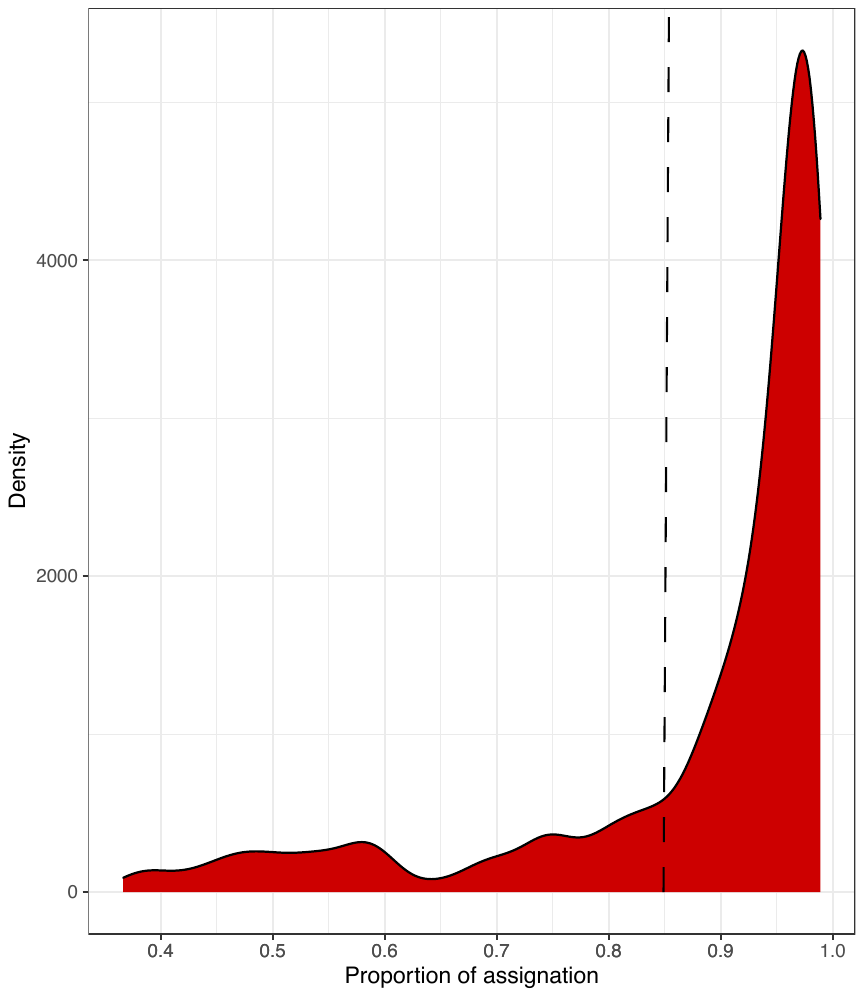

**Figure S3. Distribution of the maximum membership coefficients inferred with STRUCTURE** **for the wild and cultivated apples (*N*=550, 26 SSR markers) at *K*=12**. The vertical line at 0.85 represents the threshold used to assign an individual to a given cluster. Individuals with a membership coefficient > 0.85 were considered as admixed genotypes (in grey in other figures).

**Table S6.** Pairwise genetic differentiation estimates (*F_ST_*, lower triangle, and Jost’s *D,* upper triangle) between the thirteen populations (i.e., groups including individuals assigned with a membership coefficient > 0.85 to a given cluster) detected with STRUCTURE at *K*=12 (*N*=424, 26 microsatellite markers). All estimates were significant, except between the wild and cultivated purple populations. Note that the purple population was split into two populations: wild and cultivated purple samples.

|  |  |  | **Western (orange)** | **Central (blue)** | **Southern (brown)** | **Lorestan (light green)** | **Kurdestan (red)** | **Hyrcanian (pink)** | **Hyrcanian (purple)** | *Malus sieversii* | *Malus sieversii* | ***Malus baccata*** | ***Malus domestica*** | **Cult. Iran (purple)** | **Cult. Iran (dark green)** |
| --- | --- | --- | --- | --- | --- | --- | --- | --- | --- | --- | --- | --- | --- | --- | --- |
| wild | Armenia | Western (orange) | -- | 0.212 | 0.213 | 0.660 | 0.475 | 0.394 | 0.695 | 0.595 | 0.357 | 0.791 | 0.504 | 0.685 | 0.623 |
|  |  | Central (blue) | 0.043 | -- | 0.255 | 0.669 | 0.543 | 0.465 | 0.719 | 0.685 | 0.422 | 0.765 | 0.578 | 0.703 | 0.661 |
|  |  | Southern (brown) | 0.041 | 0.062 | -- | 0.667 | 0.541 | 0.425 | 0.785 | 0.551 | 0.348 | 0.789 | 0.602 | 0.760 | 0.663 |
|  | Iran | Lorestan (light green) | 0.22 | 0.233 | 0.246 | -- | 0.250 | 0.222 | 0.364 | 0.267 | 0.295 | 0.906 | 0.285 | 0.380 | 0.338 |
|  |  | Kurdestan (red) | 0.107 | 0.144 | 0.145 | 0.25 | -- | 0.374 | 0.663 | 0.586 | 0.574 | 0.871 | 0.715 | 0.649 | 0.627 |
|  |  | Hyrcanian (pink) | 0.082 | 0.114 | 0.105 | 0.222 | 0.109 | -- | 0.699 | 0.468 | 0.399 | 0.826 | 0.706 | 0.685 | 0.663 |
|  |  | Hyrcanian (purple) | 0.185 | 0.208 | 0.232 | 0.364 | 0.247 | 0.214 | -- | 0.780 | 0.748 | 0.922 | 0.412 | 0.025 | 0.514 |
|  | Kazakhstan | *Malus sieversii* | 0.081 | 0.113 | 0.096 | 0.295 | 0.18 | 0.114 | 0.258 | -- | 0.117 | 0.933 | 0.172 | 0.246 | 0.214 |
|  | Kyrkyzstan | *Malus sieversii* | 0.118 | 0.16 | 0.133 | 0.267 | 0.166 | 0.122 | 0.24 | 0.117 | -- | 0.850 | 0.165 | 0.259 | 0.237 |
|  | Russia | *Malus baccata* | 0.233 | 0.245 | 0.26 | 0.466 | 0.335 | 0.27 | 0.388 | 0.316 | 0.304 | -- | 0.285 | 0.399 | 0.389 |
| cultivated | Europe | *Malus domestica* | 0.097 | 0.134 | 0.138 | 0.285 | 0.19 | 0.168 | 0.14 | 0.165 | 0.172 | 0.285 | -- | 0.141 | 0.188 |
|  | Iran | Cult. Iran (purple) | 0.189 | 0.21 | 0.233 | 0.38 | 0.252 | 0.216 | 0.014 | 0.259 | 0.246 | 0.399 | 0.141 | -- | 0.544 |
|  |  | Cult. Iran (dark_green) | 0.147 | 0.183 | 0.186 | 0.338 | 0.221 | 0.193 | 0.226 | 0.237 | 0.214 | 0.389 | 0.188 | 0.246 | -- |

**Table S7.** Pairwise genetic differentiation estimates (*F_ST_*, lower triangle, and Jost’s *D,* upper triangle) between the twelve populations (*i.e.*, groups including individuals assigned with a membership coefficient > 0.85 to a given cluster) detected with STRUCTURE at *K*=12 (*N* = 424, 26 microsatellite markers). All estimates were significant, except between the wild and cultivated purple populations.

| Population | Western (orange) | Central (blue) | Southern (brown) | Lorestan (light green) | Kurdestan (red) | Hyrcanian (pink) | Hyrcanian (purple) | (light blue) | (cyan) | (light red) | European cultivars | **Iranian cultivars (purple)** | **Iranian cultivars (dark green)** |
| --- | --- | --- | --- | --- | --- | --- | --- | --- | --- | --- | --- | --- | --- |
| Western (orange) |  | **0.03** | **0.02** | **0** | **0** | **0.04** | **0** | 0.99 | **0** | 0.63 | 0.51 | **0** | **0.01** |
| Central (blue) | **0** |  | 0.79 | **0** | 0.32 | 0.96 | **0** | **0.07** | **0.02** | **0.02** | 0.13 | **0** | 0.39 |
| Southern (brown) | **0.01** | 0.92 |  | **0.01** | 0.67 | 0.63 | **0.02** | **0.06** | **0.07** | **0.04** | **0.08** | **0** | 0.65 |
| Lorestan (light green) | **0** | **0** | **0** |  | **0.02** | **0** | 0.53 | **0** | 0.35 | **0** | **0** | 0.82 | **0.04** |
| Kurdestan (red) | **0** | **0.01** | **0.01** | **0** |  | 0.32 | **0.02** | **0.01** | 0.12 | **0** | **0.01** | **0** | 0.96 |
| Hyrcanian (pink) | **0** | 0.62 | 0.56 | **0** | 0.11 |  | **0** | **0.09** | **0.02** | **0.05** | **0.20** | **0** | **0.32** |
| Hyrcanian (purple) | **0** | **0** | **0** | **0** | **0.02** | **0** |  | **0** | 0.50 | **0** | **0** | **0.08** | **0.06** |
| (light blue) | **0** | 0.59 | 0.48 | **0** | **0.04** | 0.92 | **0** |  | **0** | 0.58 | 0.67 | **0** | **0.01** |
| (cyan) | **0** | **0.01** | **0** | **0** | 0.93 | **0.09** | **0.02** | **0.05** |  | **0** | **0** | **0.07** | 0.22 |
| (light red) | **0** | **0** | **0** | **0** | **0.05** | **0** | 0.61 | **0** | **0.05** |  | 0.37 | **0** | **0.01** |
| European cultivars | **0** | 0.61 | 0.63 | **0** | **0** | 0.93 | **0** | 1.00 | **0.01** | **0** |  | **0** | **0.03** |
| **Iranian cultivars (purple)** | **0** | **0** | **0** | **0** | **0** | **0** | **0.50** | **0** | **0** | 0.99 | **0** |  | **0.01** |
| **Iranian cultivars (dark green)** | **0** | **0** | **0** | **0** | 0.16 | **0** | 0.62 | **0** | 0.13 | 0.39 | **0** | 0.20 |  |

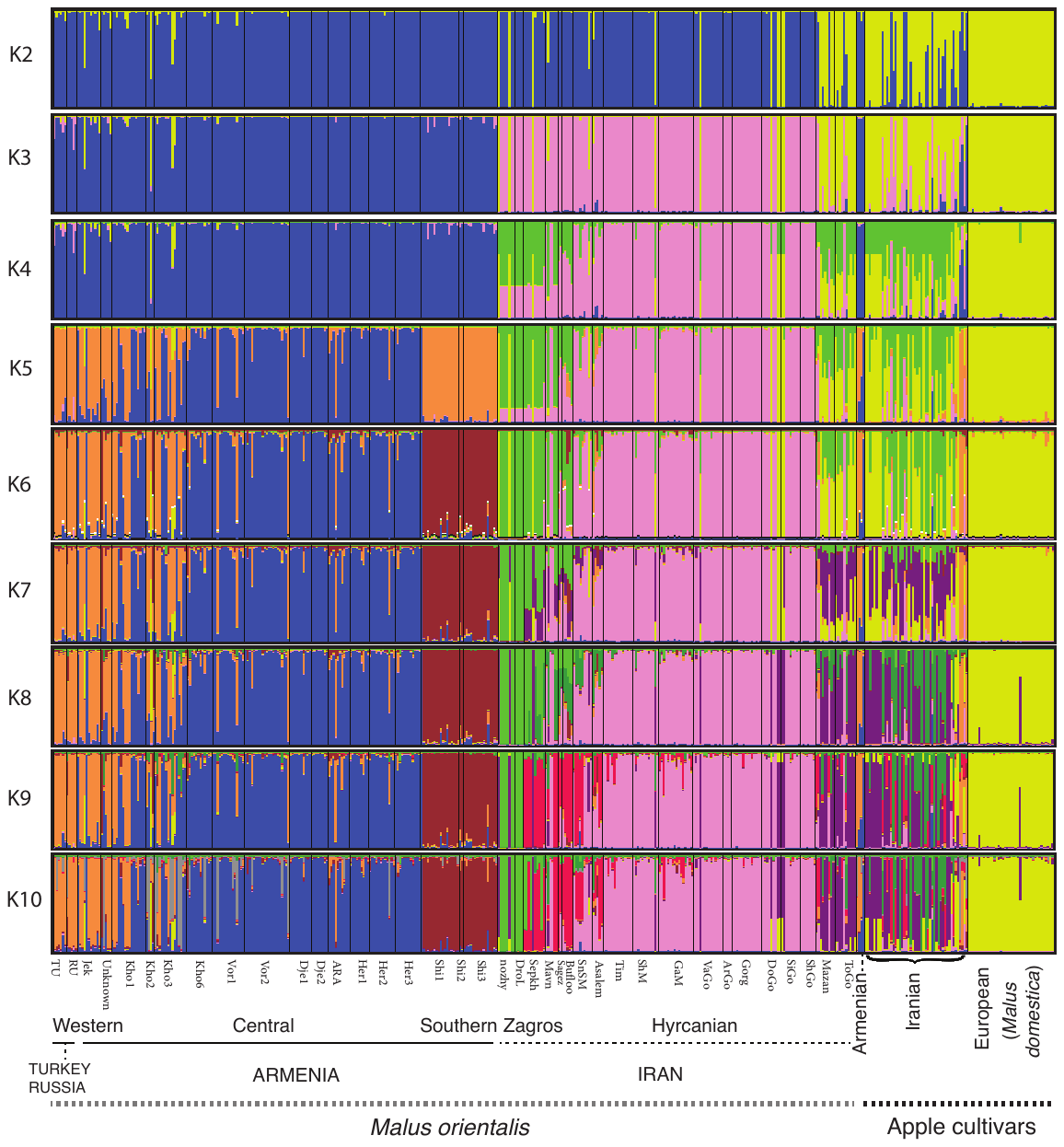

**Figure S4.** **Bayesian clustering obtained with STRUCTURE for the wild and cultivated apple using 26 microsatellite markers, from *K*=2 to *K*=10, based on the dataset excluding *M. sieversii* and *M. baccata* (*N*=466).** Each individual is represented by a vertical bar, partitioned into *K* segments representing the proportions of ancestry of its genome in *K* clusters. Armenian sites (Shi1: Shikahogh1, Shi2: Shikahogh2, Shi3: Shikahogh3, Kho1: KhosrovReserve1, Kho2: KhosrovReserve2, Kho3: KhosrovReserve3, Kho3bis: KhosrovReserve3bis; Kho3ter: KhosrovReserve3ter, Kho4: KhosrovReserve4, Kho5: KhosrovReserve5, Kho6: KhosrovReserve6, Vor2: Vorotanpass2, Vor1: Vorotanpass1, Dje2: Djermuk2, Dje1: Djermuk1, Her2: Hermon2, Her1: Hermon1, Her3: Hermon3, ARA: ARA, Jek: Jermouck, Arm_unknown: Arm_unknown), Iranian sites (nozhy: Nozhyan region , DroL: Drood region, Sepkh: Sepedkoh region, Mavn: Marivan region, Sagez: Sagez region, Bufloo: Bufloo region, Asalem: Asalem region, Tim: Tim region; Tilak: Tilak region, SnSM: Sangdeh region, ShM: Shit region, VaM: Vaz region, GaM: Ganasara region, Mazan: Mazan region; Amol: Amol abesk region , VaGo: Vatna region, ArGo: Arsam region, Gorg: Gorgan province, DoGo: Dorak region, ToGo: Tokestan region, SiGo: Siamarzkoh region, Sh6sh: Sheshab region). The main regions in the Caucasus and/or defined in the main text are also added: Western Caucasus (Turkey and Russia), Central and Southern Armenia, Zagros and the Hyrcanian forest.

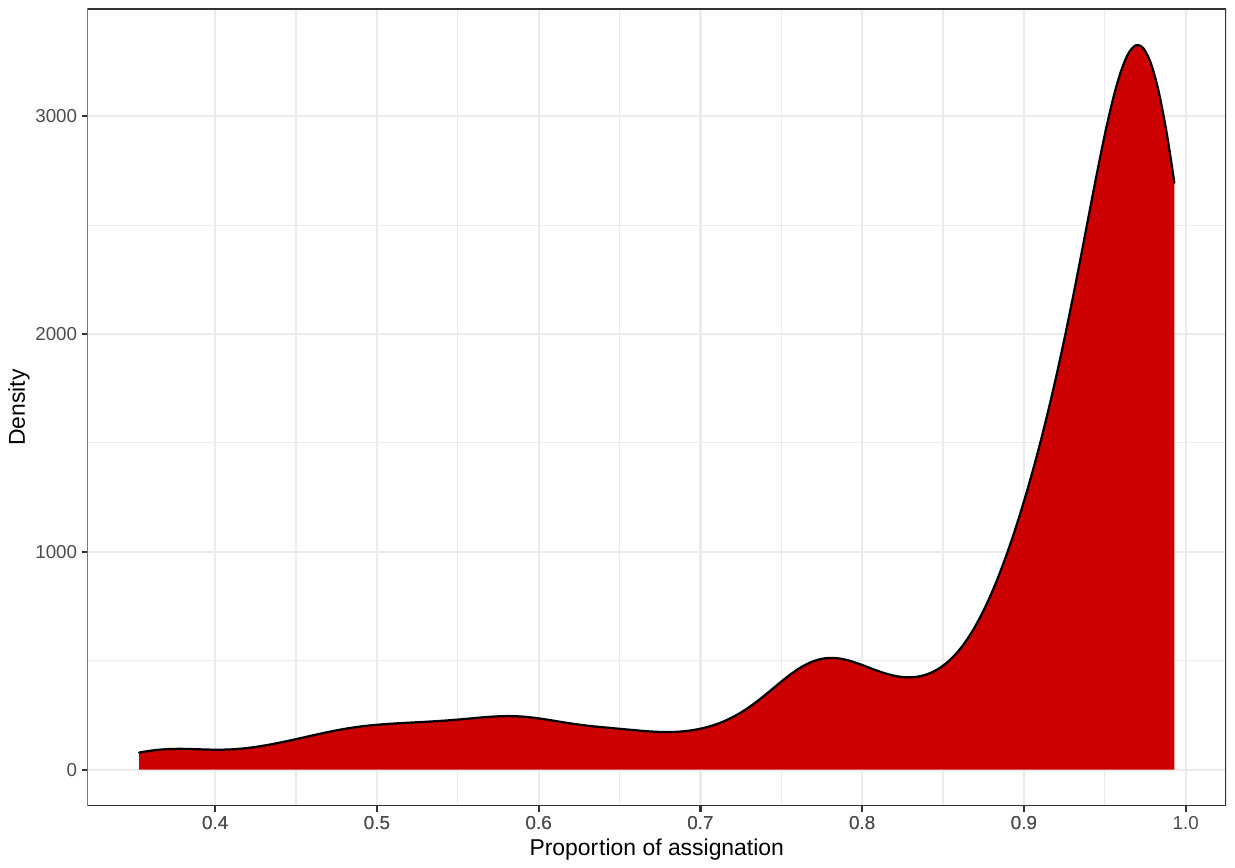

**Fig. S5. Distribution of the maximum membership coefficient inferred with STRUCTURE** **for the 446 wild and cultivated apples at *K*=9**. Individuals with a membership coefficient < 0.90 were considered as hybrids.

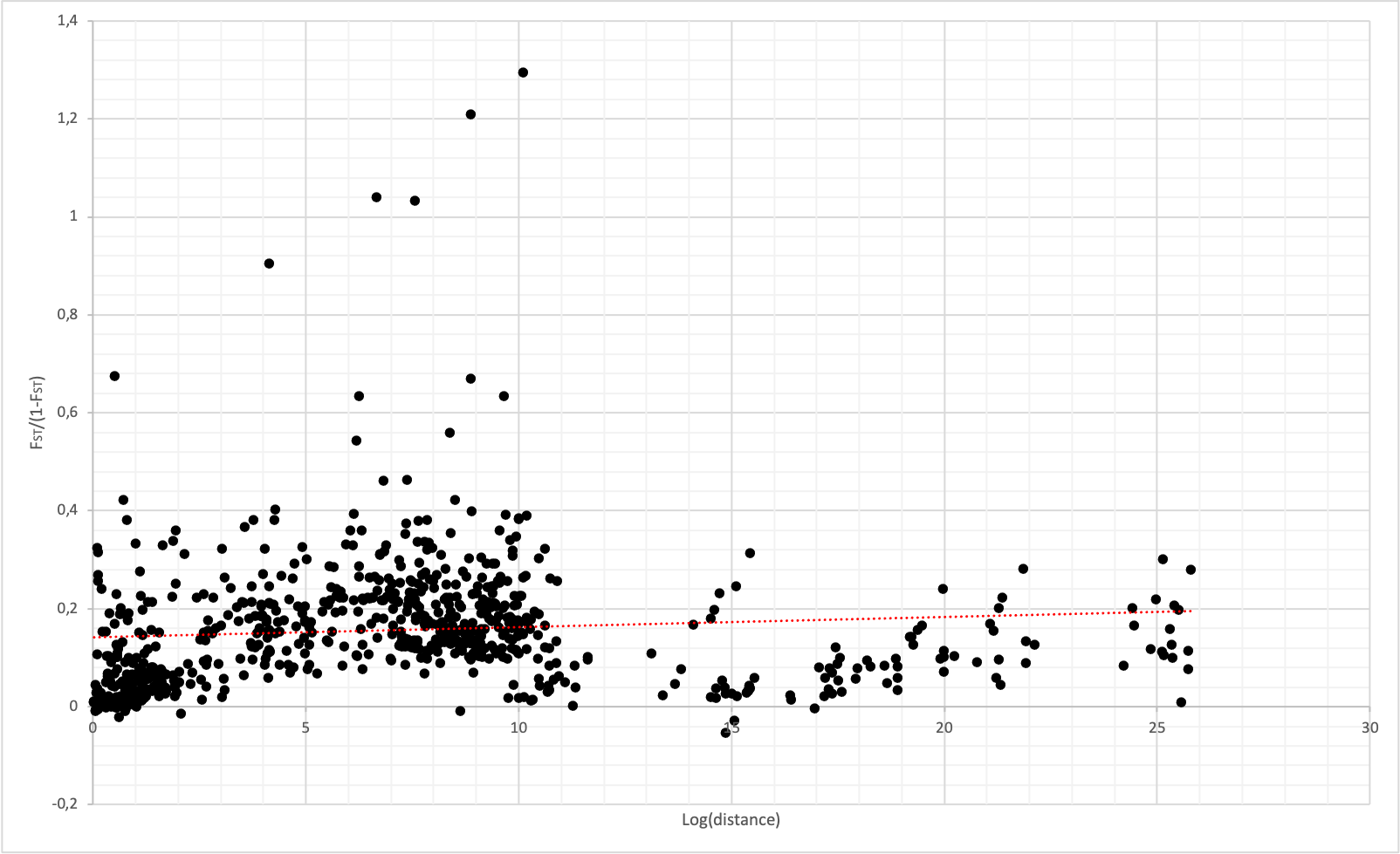

**Figure S6. Pairwise genetic differentiation (*F_ST_/(1- F_ST_)*) in function of the logarithm of the distance for *Malus orientalis* (*N*=339, 36 sample sites with at least four individuals) in the Caucasus.** The red dotted line represents the linear regression line: the adjusted R-squared and associated p-value are presented in the main text.

**Table S8.** **Matrix of gene flow inferred with BayesAss** **between the ten apple populations (*i.e.*, groups including individuals assigned with a membership coefficient > 0.90 to a given cluster, the wild and crop purple populations were split) detected with STRUCTURE at *K*=9 (*N* = 423, 26 microsatellite markers) in the Caucasus and Iran.**

|  |  | Wild (j) | | | | | | | Crop (j) | | |
| --- | --- | --- | --- | --- | --- | --- | --- | --- | --- | --- | --- |
|  |  | Western (orange) | Central (blue) | Southern (brown) | Lorestan (light green) | Kurdestan (red) | Hyrcanian (pink) | Hyrcanian (purple) | **Iranian cultivars (purple)** | **Iranian cultivars (dark green)** | European cultivars |
| Wild (i) | Western (orange) | **0.861** | 0.007 | 0.006 | 0.007 | 0.008 | 0.007 | 0.081 | 0.009 | 0.006 | 0.007 |
|  | Central (blue) | 0.017 | **0.956** | 0.010 | 0.003 | 0.002 | 0.002 | 0.003 | 0.002 | 0.003 | 0.003 |
|  | Southern (brown) | 0.006 | 0.017 | **0.920** | 0.015 | 0.008 | 0.008 | 0.009 | 0.006 | 0.006 | 0.007 |
|  | Lorestan (light green) | 0.015 | 0.020 | 0.026 | **0.830** | 0.032 | 0.015 | 0.014 | 0.018 | 0.015 | 0.017 |
|  | Kurdestan (red) | 0.010 | 0.017 | 0.025 | 0.021 | **0.877** | 0.011 | 0.010 | 0.010 | 0.010 | 0.009 |
|  | Hyrcanian (pink) | 0.003 | 0.003 | 0.002 | 0.003 | 0.014 | **0.963** | 0.003 | 0.004 | 0.003 | 0.002 |
|  | Hyrcanian (purple) | 0.010 | 0.009 | 0.006 | 0.007 | 0.009 | 0.041 | **0.897** | 0.008 | 0.007 | 0.007 |
| Crop (i) | **Iranian cultivars (purple)** | 0.013 | 0.012 | 0.014 | 0.014 | 0.012 | 0.041 | 0.076 | **0.790** | 0.012 | 0.016 |
|  | **Iranian cultivars (dark green)** | 0.014 | 0.016 | 0.014 | 0.018 | 0.019 | 0.016 | 0.020 | 0.020 | **0.846** | 0.016 |
|  | European cultivars | 0.010 | 0.009 | 0.011 | 0.006 | 0.007 | 0.009 | 0.008 | 0.009 | 0.016 | **0.916** |

Values in the form *m_ij_* represent the proportion of individuals in the *i^th^* population (crop_i_ or wild_i_) that originated from the *j^th^* population per generation. The mean posterior probability distributions were averaged over three replicate runs with different seed numbers.

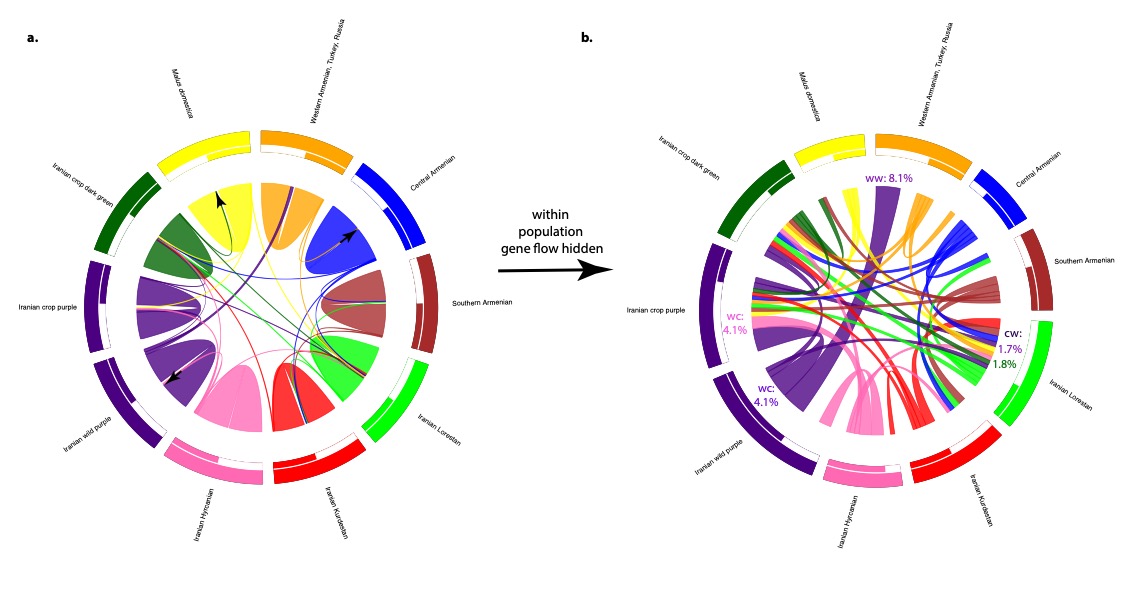

**Figure S7. Circle migration plots representing gene flow estimates among apple populations in the Caucasus and Iran inferred with BayesAss. a.** Circle migration plot with the gene flow within population (in bold in Table S8). **b.** Circle migration plot hiding gene flow within population for a better visualization of between-population gene flow. Populations are given on the outside of the circle, where each can be assumed as source population from which individuals migrated. The thickness of the arcs represents the rate of migration. *wc*, *cw* or *ww*, with their respective colors, indicated examples of wild-to-crop, crop-to-wild and wild-to-wild gene flow from Table S8.

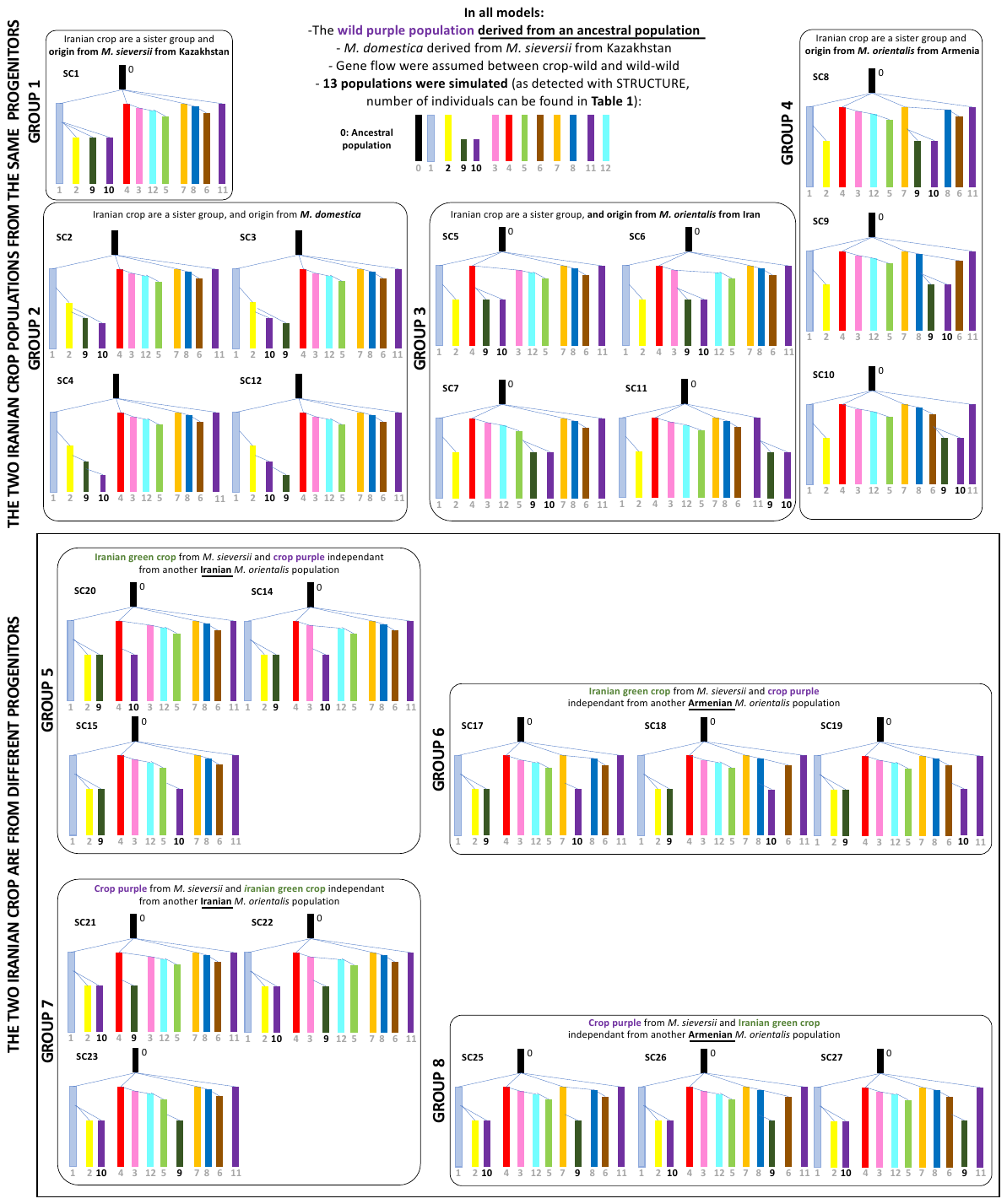

**Figure S8**. **Details of the scenarios tested with approximate Bayesian computation for reconstructing the domestication history of the Iranian cultivated apple*.*** Eight groups of scenarios were simulated assuming that the two Iranian cultivated apple populations diverged from the same (groups 1, 3 and 4) or different wild (*M. sieversii* from Kazakhstan or *M. orientalis* from Iran and Armenia, groups 5 to 8) or cultivated (*M. domestica,* group 2) apple populations*.* In all models, the wild purple population was assumed to diverge from an ancestral population (referred to as 0). A total of 24 scenarios were simulated. Bidirectional gene flow between populations were assumed (see Table S3 for priors). The different colors represent the 12 populations (+ one ancestral population in black) inferred with STRUCTURE (Figure 1, Table 1, excluding *M. baccata*). Each population is represented by a number as depicted in the upper part of the figure. Scenario identification number (SC*X*) is arbitrary.

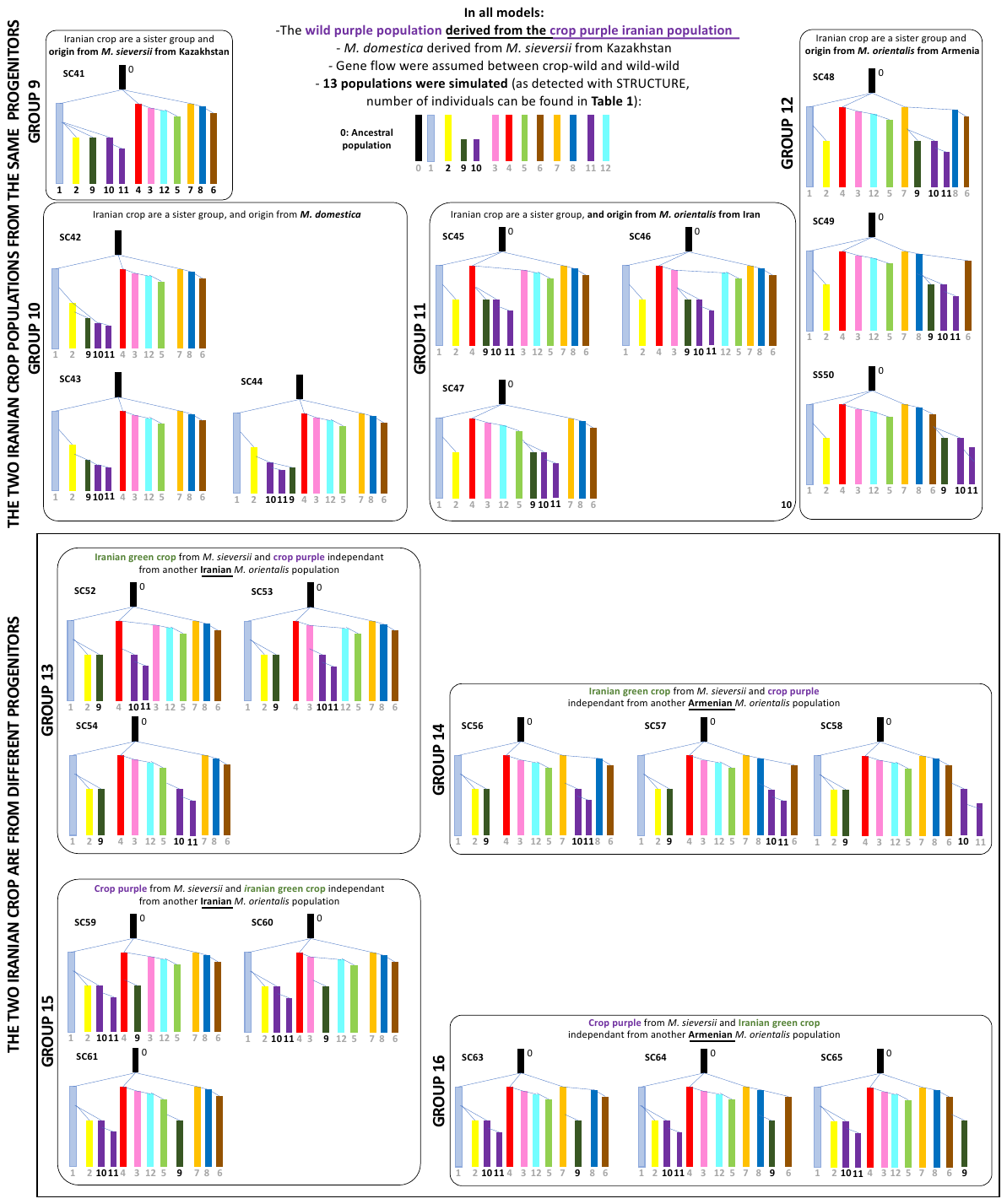
**Figure S9**. **Details of the scenarios tested with approximate Bayesian computation for reconstructing the domestication history of the Iranian cultivated apple*.*** Eight groups of scenarios were simulated assuming that the two Iranian cultivated apple populations diverged from the same (groups 9,11 and 12) or different (groups 13 to 16) wild (*M. sieversii* from Kazakhstan or *M. orientalis* from Iran and Armenia) or cultivated (*M. domestica*, group 10) apple populations*.* In all models, the wild purple population was assumed to diverge from the crop purple population (referred as to “0”). A total of 22 scenarios were simulated. Bidirectional gene flow between populations were assumed (see Table S3 for priors). The different colors represent the 12 populations (+ one ancestral in black) inferred with STRUCTURE (Figure 1, Table 1, excluding *M. baccata*). Each population is represented by a number as depicted in the upper part of the figure. Scenario identification number (SC*X*) is arbitrary.

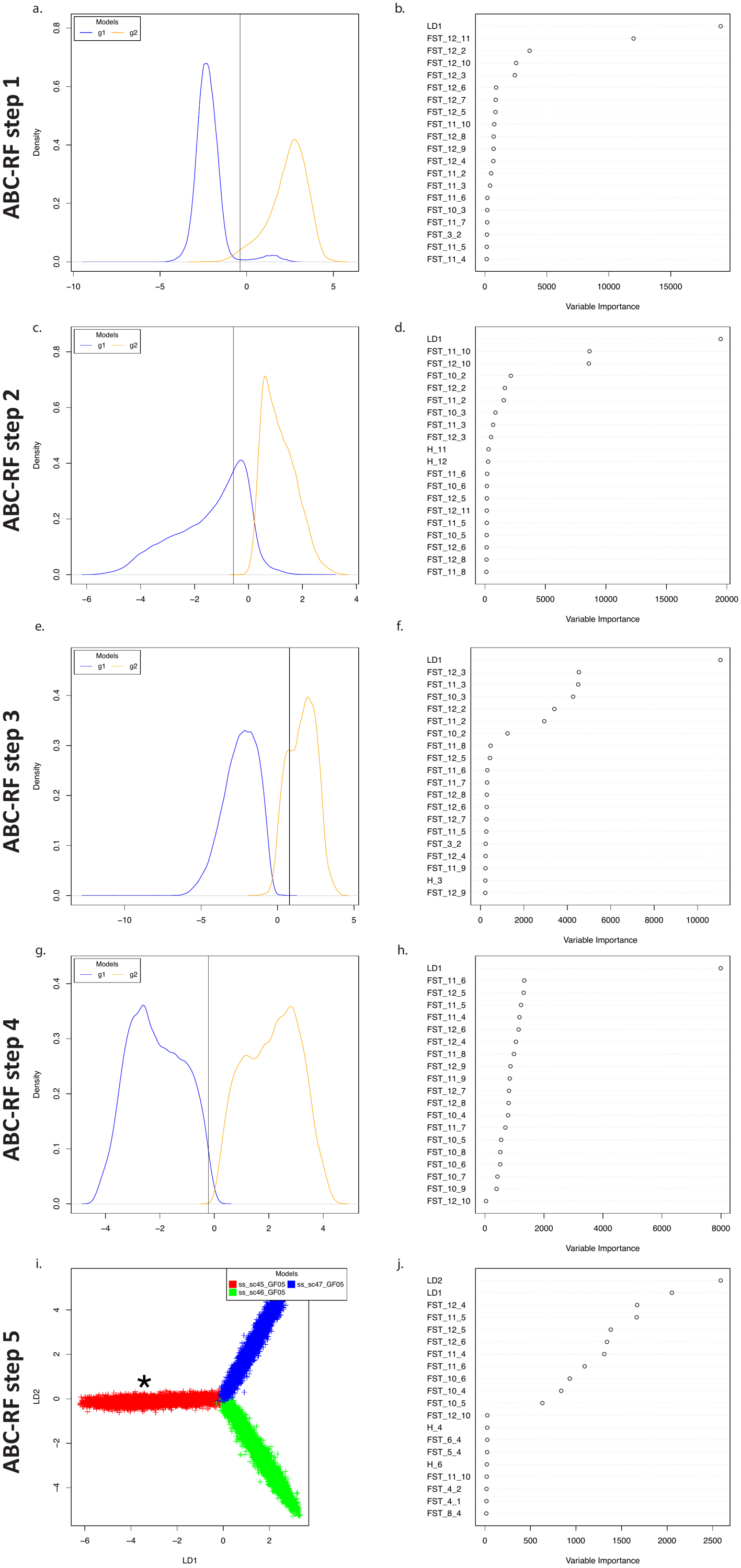

### **Figure. S10.** Linear discriminant analysis, and associated variance plot, obtained for each of the five random forest approximate Bayesian computation steps to infer the Iranian cultivated evolutionary history.

**Table S9. Results of the ABC-RF algorithm inferring the origin of the wild Iranian apple population (ABC-RF step 1).** Scenarios assumed gene flow among wild and crop populations. We reported in this table: repartition of votes among the two groups of scenarios for each replicate, and mean and standard deviations over replicates for each group of scenarios, posterior probability and prior error rate for the best scenario, *i.e.*, the scenario with the highest number of votes, respectively. The most likely model is highlighted in bold (10 out of 10 votes for the scenario assuming the wild and cultivated apple purple populations as a sister group).

| Replicates | Groups 1 to 8 | **Groups 9 to 16** | Posterior probability % | Prior error rate |
| --- | --- | --- | --- | --- |
| 1 | 214 | **286** | 63 | 0.98 |
| 2 | 231 | **269** | 68 | 0.98 |
| 3 | 208 | **292** | 66 | 0.98 |
| 4 | 221 | **279** | 64 | 0.98 |
| 5 | 221 | **279** | 68 | 0.98 |
| 6 | 195 | **305** | 65 | 0.98 |
| 7 | 229 | **271** | 67 | 0.98 |
| 8 | 225 | **275** | 64 | 0.99 |
| 9 | 213 | **287** | 61 | 0.98 |
| 10 | 224 | **276** | 67 | 0.99 |
| Mean | 218.1 | **281.9** | **65** | **0.98** |
| sd | 10.9 | **10.9** | 2 | 0.00 |

**Table S10. Results of the ABC-RF algorithm inferring whether the two cultivated Iranian populations are a sister group which diverged from a same wild population (groups 9 to 12) or diverged from two different wild populations (groups 13 to 16, Figure S8) (ABC-RF step 2).** Scenarios assumed gene flow among wild and crop populations. We reported in this table: repartition of votes among the two groups of scenarios for each replicate, and mean and standard deviations over replicates for each group of scenarios, posterior probability and prior error rate for the best scenario, *i.e.*, the scenario with the highest number of votes, respectively. The most likely model is highlighted in bold (10 out of 10 votes for the scenario assuming that the Iranian cultivated apple populations are a sister group).

| **Replicates** | **Groups 9 to 12** | **Groups 13 to 16** | **Posterior probability (%)** | **Prior error rate** |
| --- | --- | --- | --- | --- |
| 1 | **295** | 205 | 80 | 1.10 |
| 2 | **313** | 187 | 83 | 1.09 |
| 3 | **287** | 213 | 82 | 1.09 |
| 4 | **291** | 209 | 82 | 1.09 |
| 5 | **299** | 201 | 80 | 1.08 |
| 6 | **292** | 208 | 84 | 1.09 |
| 7 | **298** | 202 | 80 | 1.10 |
| 8 | **288** | 212 | 82 | 1.10 |
| 9 | **296** | 204 | 85 | 1.08 |
| 10 | **300** | 200 | 80 | 1.07 |
| Mean | **296** | 204 | 82 | 1.09 |
| Standard deviation | **7.5** | 7.5 | 2 | 0.01 |

**Table S11. Results of the ABC-RF algorithm inferring whether sister cultivated apple populations originated from *M. sieversii/M.domestica* (groups 9 and 10) or *M. orientalis* (groups 11 and 12) (ABC-RF step 3).** Scenarios assumed gene flow among wild and crop populations. We reported in this table: repartition of votes among the two groups of scenarios for each replicate, and mean and standard deviations over replicates for each group of scenarios, posterior probability and prior error rate for the best scenario, *i.e.*, the scenario with the highest number of votes, respectively. The most likely model is highlighted in bold (10 out of 10 votes for the scenario assuming an origin of the sister Iranian cultivated apple populations from *M. orientalis*).

| **Replicates** | **Groups 9+10** | **Groups 11+12** | **Posterior probability (%)** | **Prior error rate** |
| --- | --- | --- | --- | --- |
| 1 | 208 | **292** | 86 | 0.20 |
| 2 | 206 | **294** | 83 | 0.21 |
| 3 | 195 | **305** | 78 | 0.22 |
| 4 | 216 | **284** | 76 | 0.21 |
| 5 | 213 | **287** | 77 | 0.20 |
| 6 | 211 | **289** | 80 | 0.21 |
| 7 | 197 | **303** | 76 | 0.22 |
| 8 | 212 | **288** | 78 | 0.22 |
| 9 | 220 | **280** | 76 | 0.21 |
| 10 | 218 | **282** | 78 | 0.21 |
| Mean | 210 | **290** | **79** | **0.21** |
| Standard deviation | 8.3 | **8.3** | 3 | 0.01 |

**Table S12. Results of the ABC-RF algorithm infering whether sister cultivated apple populations originated from *M. orientalis* from Iran (group 11) or from Armenia (group 12) (ABC-RF step 4).** Scenarios assumed gene flow among wild and crop populations. We reported in this table: repartition of votes among the two groups of scenarios for each replicate, and mean and standard deviations over replicates for each group of scenarios, posterior probability and prior error rate for the best scenario, *i.e.*, the scenario with the highest number of votes, respectively. The most likely model is highlighted in bold (10 out of 10 votes for the scenario assuming an origin of the sister Iranian cultivated apple populations from *M. orientalis* from Iran).

| **Replicates** | **Group 11** | **Group 12** | **Posterior probability (%)** | **Prior error rate** |
| --- | --- | --- | --- | --- |
| 1 | **324** | 176 | 89 | 0.10 |
| 2 | **314** | 186 | 96 | 0.10 |
| 3 | **323** | 177 | 90 | 0.09 |
| 4 | **301** | 199 | 91 | 0.10 |
| 5 | **311** | 189 | 92 | 0.10 |
| 6 | **313** | 187 | 91 | 0.10 |
| 7 | **313** | 187 | 91 | 0.10 |
| 8 | **315** | 185 | 91 | 0.09 |
| 9 | **307** | 193 | 91 | 0.10 |
| 10 | **339** | 161 | 92 | 0.09 |
| Mean | **316** | 184 | **92** | **0.10** |
| Standard deviation | **10.52** | 10.52 | 2 | 0 |

**Table S13. Results of the ABC-RF algorithm inferring whether the sister Iranian cultivated apple populations originated from one of the three Iranian *M. orientalis* populations (ABC-RF step 5).** Scenarios assumed gene flow among wild and crop populations. We reported in this table: repartition of votes among the two groups of scenarios for each replicate, and mean and standard deviations over replicates for each group of scenarios, posterior probability and prior error rate for the best scenario, *i.e.*, the scenario with the highest number of votes, respectively. The most likely model is highlighted in bold (10 out of 10 votes for the scenario assuming an origin of the sister Iranian cultivated apple populations from the Iranian red *M. orientalis* population).

| **Replicates** | **sc45** | **sc46** | **sc47** | **Posterior probability (%)** | **Prior error rate** |
| --- | --- | --- | --- | --- | --- |
| 1 | **253** | 79 | 168 | 90 | 0.11 |
| 2 | **267** | 68 | 165 | 89 | 0.11 |
| 3 | **240** | 93 | 167 | 93 | 0.11 |
| 4 | **245** | 71 | 184 | 88 | 0.10 |
| 5 | **274** | 75 | 151 | 93 | 0.10 |
| 6 | **229** | 91 | 180 | 92 | 0.12 |
| 7 | **230** | 81 | 189 | 89 | 0.12 |
| 8 | **241** | 86 | 173 | 92 | 0.11 |
| 9 | **238** | 73 | 189 | 90 | 0.11 |
| 10 | **242** | 89 | 169 | 88 | 0.11 |
| Mean | **245.9** | 80.6 | 173.5 | 90 | 0.11 |
| Standard deviation | **14.76** | 8.85 | 12.04 | 2 | 0.01 |

**Table S14. Parameter estimates infered for the Iranian cultivated apple domestication history, computed with the most likely (SC45, Figure S8).** We computed the median and mean posterior probability, the 90% confidence interval (q5% and q95%) and the NMAE which significates the facility of estimating the associate parameter. Divergence time estimates were multiplied by 10, which is the generation time estimates for apple, to transform them in years. Parameters are described in Table S3.

| Parameters | Median | q5% | q95% | NMAE |
| --- | --- | --- | --- | --- |
| *N_ANC_* | 123 | 57 | 197 | 0.30 |
| *N_blue_* | 82 | 53 | 99 | 0.17 |
| *N_brown_* | 79 | 53 | 100 | 0.17 |
| *N_crop_ir_purple_* | 73 | 50 | 98 | 0.18 |
| *N_cyan_* | 77 | 50 | 99 | 0.16 |
| *N_crop_dark_green_* | 76 | 51 | 100 | 0.17 |
| *N_light_blue_* | 71 | 50 | 99 | 0.18 |
| *N_light_green_* | 72 | 51 | 99 | 0.17 |
| *N_orange_* | 75 | 50 | 99 | 0.16 |
| *N_pink_* | 75 | 52 | 99 | 0.16 |
| *N_red_* | 74 | 51 | 98 | 0.17 |
| *N_wild_ir_purple_* | 72 | 51 | 99 | 0.17 |
| *N_crop_domestica_* | 76 | 51 | 99 | 0.17 |
| *μ* | 0.00082 | 0.00038 | 0.00098 | 0.25 |
| *T_WILD_brown_* | 258462 | 35260 | 381050 | 0.98 |
| *T _WILD_light_green_* | 212720 | 37110 | 375840 | 0.99 |
| *T _WILD_purple_* | 418 | 100 | 2500 | 0.67 |
| *T_CROP_ir_green_* | 836 | 130 | 5030 | 0.71 |
| *T_CROP_ir_purple_* | 803 | 160 | 3969 | 0.73 |
| *T_CROP_domestica_* | 3898 | 270 | 7670 | 0.54 |
| *T_WILD_siev_* | 194130 | 37740 | 388286 | 1.04 |
| *T_WILD_blue_* | 311902 | 109440 | 599780 | 1 |
| *T_WILD_lightblue_* | 332695 | 77480 | 669916 | 1 |
| *T_WILD_orange_* | 501804 | 189470 | 882770 | 0 |
| *T_WILD_pink_* | 453580 | 104160 | 921823 | 0 |
| *T_WILD_red_* | 637090 | 216217 | 1210480 | 0 |
| *m_10_1* | 6.86E-05 | 1.24E-06 | 9.34E-05 | 4.37 |
| *m_10_11* | 4.52E-05 | 1.55E-06 | 9.06E-05 | 4.12 |
| *m_10_12* | 5.79E-05 | 1.30E-06 | 9.50E-05 | 4.26 |
| *m_10_2* | 4.72E-05 | 1.37E-06 | 8.94E-05 | 4.45 |
| *m_10_3* | 6.34E-05 | 1.15E-06 | 9.56E-05 | 4.39 |
| *m_10_4* | 6.91E-05 | 1.73E-06 | 9.47E-05 | 4.61 |
| *m_10_5* | 5.85E-05 | 1.32E-06 | 9.87E-05 | 4.22 |
| *m_10_6* | 6.72E-05 | 1.64E-06 | 9.45E-05 | 4.22 |
| *m_10_7* | 5.13E-05 | 1.70E-06 | 9.90E-05 | 4.17 |
| *m_10_8* | 6.78E-05 | 2.17E-06 | 9.75E-05 | 4.30 |
| *m_10_9* | 5.69E-05 | 1.26E-06 | 9.11E-05 | 3.70 |
| *m_11_1* | 5.23E-05 | 1.56E-06 | 9.65E-05 | 3.99 |
| *m_11_10* | 6.13E-05 | 1.31E-06 | 9.70E-05 | 3.55 |
| *m_11_12* | 5.71E-05 | 1.50E-06 | 9.77E-05 | 4.20 |
| *m_11_2* | 7.12E-05 | 1.46E-06 | 9.74E-05 | 4.08 |
| *m_11_3* | 5.10E-05 | 1.36E-06 | 9.67E-05 | 4.49 |
| *m_11_4* | 5.11E-05 | 1.73E-06 | 9.67E-05 | 4.15 |
| *m_11_5* | 5.54E-05 | 1.33E-06 | 9.66E-05 | 4.47 |
| *m_11_6* | 6.11E-05 | 1.03E-06 | 9.12E-05 | 3.99 |
| *m_11_7* | 5.43E-05 | 2.03E-06 | 9.81E-05 | 4.19 |
| *m_11_8* | 6.60E-05 | 2.75E-06 | 9.99E-05 | 4.00 |
| *m_11_9* | 5.90E-05 | 1.51E-06 | 9.64E-05 | 4.07 |
| *m_12_1* | 5.24E-05 | 1.71E-06 | 9.76E-05 | 4.26 |
| *m_12_10* | 4.79E-05 | 1.82E-06 | 9.50E-05 | 4.15 |
| *m_12_11* | 3.85E-05 | 1.19E-06 | 9.90E-05 | 4.04 |
| *m_12_2* | 6.30E-05 | 2.26E-06 | 8.87E-05 | 4.45 |
| *m_12_3* | 5.84E-05 | 1.47E-06 | 9.83E-05 | 3.97 |
| *m_12_4* | 5.11E-05 | 1.59E-06 | 9.99E-05 | 4.08 |
| *m_12_5* | 6.81E-05 | 1.48E-06 | 9.77E-05 | 4.25 |
| *m_12_6* | 6.38E-05 | 1.78E-06 | 9.74E-05 | 4.28 |
| *m_12_7* | 5.70E-05 | 2.00E-06 | 9.97E-05 | 4.46 |
| *m_12_8* | 6.10E-05 | 1.36E-06 | 9.73E-05 | 4.29 |
| *m_12_9* | 6.07E-05 | 1.70E-06 | 9.14E-05 | 4.13 |
| *m_1_10* | 4.88E-05 | 1.29E-06 | 9.54E-05 | 4.37 |
| *m_1_11* | 6.12E-05 | 1.65E-06 | 9.96E-05 | 3.91 |
| *m_1_12* | 6.60E-05 | 1.81E-06 | 9.97E-05 | 4.52 |
| *m_1_2* | 7.11E-05 | 2.06E-06 | 9.89E-05 | 4.16 |
| *m_1_3* | 5.20E-05 | 1.60E-06 | 9.58E-05 | 3.97 |
| *m_1_4* | 5.70E-05 | 1.45E-06 | 9.34E-05 | 4.13 |
| *m_1_5* | 5.03E-05 | 1.67E-06 | 9.98E-05 | 4.22 |
| *m_1_6* | 5.22E-05 | 1.39E-06 | 9.80E-05 | 4.30 |
| *m_1_7* | 6.37E-05 | 1.33E-06 | 9.26E-05 | 4.05 |
| *m_1_8* | 5.01E-05 | 1.33E-06 | 9.31E-05 | 4.35 |
| *m_1_9* | 6.97E-05 | 1.46E-06 | 9.89E-05 | 4.04 |
| *m_2_1* | 3.11E-05 | 1.24E-06 | 9.44E-05 | 4.33 |
| *m_2_10* | 7.20E-05 | 2.85E-06 | 9.73E-05 | 4.71 |
| *m_2_11* | 4.87E-05 | 2.59E-06 | 9.80E-05 | 3.77 |
| *m_2_12* | 7.02E-05 | 1.48E-06 | 9.93E-05 | 4.32 |
| *m_2_3* | 7.19E-05 | 1.47E-06 | 9.77E-05 | 4.46 |
| *m_2_4* | 5.43E-05 | 1.88E-06 | 9.94E-05 | 4.15 |
| *m_2_5* | 4.81E-05 | 1.48E-06 | 9.48E-05 | 4.33 |
| *m_2_6* | 4.14E-05 | 1.97E-06 | 9.93E-05 | 4.39 |
| *m_2_7* | 5.51E-05 | 1.80E-06 | 9.98E-05 | 4.56 |
| *m_2_8* | 5.78E-05 | 1.82E-06 | 9.33E-05 | 4.37 |
| *m_2_9* | 8.82E-05 | 1.67E-06 | 9.91E-05 | 3.85 |
| *m_3_1* | 5.83E-05 | 1.34E-06 | 8.35E-05 | 4.06 |
| *m_3_10* | 5.44E-05 | 1.17E-06 | 9.86E-05 | 4.11 |
| *m_3_11* | 3.43E-05 | 2.04E-06 | 9.86E-05 | 4.35 |
| *m_3_12* | 5.70E-05 | 1.42E-06 | 9.76E-05 | 4.37 |
| *m_3_2* | 6.54E-05 | 1.74E-06 | 9.87E-05 | 4.17 |
| *m_3_4* | 5.40E-05 | 1.14E-06 | 9.44E-05 | 4.03 |
| *m_3_5* | 5.80E-05 | 1.67E-06 | 9.91E-05 | 4.51 |
| *m_3_6* | 7.88E-05 | 1.43E-06 | 9.69E-05 | 4.26 |
| *m_3_7* | 4.85E-05 | 1.36E-06 | 9.43E-05 | 4.52 |
| *m_3_8* | 5.68E-05 | 1.14E-06 | 9.90E-05 | 4.37 |
| *m_3_9* | 8.39E-05 | 1.51E-06 | 9.80E-05 | 4.22 |
| *m_4_1* | 6.24E-05 | 1.50E-06 | 9.24E-05 | 4.41 |
| *m_4_10* | 7.82E-05 | 1.80E-06 | 9.78E-05 | 4.42 |
| *m_4_11* | 5.33E-05 | 1.26E-06 | 9.78E-05 | 4.46 |
| *m_4_12* | 5.10E-05 | 1.25E-06 | 9.98E-05 | 4.71 |
| *m_4_2* | 8.43E-05 | 2.57E-06 | 9.68E-05 | 4.10 |
| *m_4_3* | 4.99E-05 | 2.41E-06 | 9.78E-05 | 4.28 |
| *m_4_5* | 7.17E-05 | 1.70E-06 | 9.56E-05 | 3.88 |
| *m_4_6* | 4.81E-05 | 1.29E-06 | 9.48E-05 | 4.11 |
| *m_4_7* | 4.48E-05 | 1.63E-06 | 9.70E-05 | 4.06 |
| *m_4_8* | 5.07E-05 | 1.37E-06 | 9.55E-05 | 4.43 |
| *m_4_9* | 4.42E-05 | 1.15E-06 | 9.52E-05 | 4.64 |
| *m_5_1* | 5.42E-05 | 1.54E-06 | 9.13E-05 | 3.94 |
| *m_5_10* | 5.27E-05 | 1.33E-06 | 9.31E-05 | 4.38 |
| *m_5_11* | 5.80E-05 | 1.72E-06 | 9.80E-05 | 4.08 |
| *m_5_12* | 6.80E-05 | 1.57E-06 | 9.38E-05 | 4.29 |
| *m_5_2* | 5.85E-05 | 1.64E-06 | 9.84E-05 | 3.94 |
| *m_5_3* | 6.63E-05 | 1.37E-06 | 9.76E-05 | 4.19 |
| *m_5_4* | 7.10E-05 | 2.02E-06 | 9.94E-05 | 4.47 |
| *m_5_6* | 7.75E-05 | 2.10E-06 | 9.02E-05 | 4.18 |
| *m_5_7* | 7.59E-05 | 1.71E-06 | 9.76E-05 | 4.23 |
| *m_5_8* | 5.48E-05 | 1.48E-06 | 9.26E-05 | 4.32 |
| *m_5_9* | 6.93E-05 | 1.61E-06 | 9.77E-05 | 4.54 |
| *m_6_1* | 6.64E-05 | 1.34E-06 | 9.69E-05 | 4.21 |
| *m_6_10* | 4.86E-05 | 1.24E-06 | 9.33E-05 | 4.37 |
| *m_6_11* | 4.86E-05 | 1.61E-06 | 9.87E-05 | 4.40 |
| *m_6_12* | 6.11E-05 | 1.48E-06 | 9.91E-05 | 4.21 |
| *m_6_2* | 6.60E-05 | 1.40E-06 | 9.48E-05 | 4.21 |
| *m_6_3* | 7.55E-05 | 1.12E-06 | 9.86E-05 | 4.62 |
| *m_6_4* | 5.95E-05 | 1.40E-06 | 9.68E-05 | 4.34 |
| *m_6_5* | 4.96E-05 | 1.86E-06 | 9.39E-05 | 4.11 |
| *m_6_7* | 6.73E-05 | 1.33E-06 | 9.46E-05 | 4.24 |
| *m_6_8* | 5.69E-05 | 1.26E-06 | 9.32E-05 | 3.86 |
| *m_6_9* | 6.13E-05 | 1.37E-06 | 9.64E-05 | 4.58 |
| *m_7_1* | 5.40E-05 | 1.01E-06 | 8.68E-05 | 4.54 |
| *m_7_10* | 7.13E-05 | 2.06E-06 | 9.70E-05 | 4.26 |
| *m_7_11* | 4.66E-05 | 1.40E-06 | 9.77E-05 | 4.53 |
| *m_7_12* | 6.04E-05 | 1.80E-06 | 9.51E-05 | 4.42 |
| *m_7_2* | 5.01E-05 | 1.35E-06 | 8.77E-05 | 4.14 |
| *m_7_3* | 6.71E-05 | 1.27E-06 | 9.80E-05 | 4.40 |
| *m_7_4* | 4.92E-05 | 1.51E-06 | 9.82E-05 | 4.04 |
| *m_7_5* | 7.04E-05 | 1.86E-06 | 9.91E-05 | 4.18 |
| *m_7_6* | 6.17E-05 | 1.24E-06 | 9.56E-05 | 4.33 |
| *m_7_8* | 6.82E-05 | 1.85E-06 | 9.91E-05 | 4.02 |
| *m_7_9* | 4.96E-05 | 1.67E-06 | 9.40E-05 | 4.47 |
| *m_8_1* | 5.70E-05 | 2.46E-06 | 9.34E-05 | 4.35 |
| *m_8_10* | 7.43E-05 | 2.10E-06 | 9.61E-05 | 3.82 |
| *m_8_11* | 4.52E-05 | 2.02E-06 | 9.80E-05 | 4.35 |
| *m_8_12* | 5.43E-05 | 1.01E-06 | 9.36E-05 | 4.41 |
| *m_8_2* | 6.69E-05 | 1.30E-06 | 9.83E-05 | 4.56 |
| *m_8_3* | 5.65E-05 | 1.67E-06 | 9.67E-05 | 4.52 |
| *m_8_4* | 5.97E-05 | 1.72E-06 | 9.65E-05 | 4.01 |
| *m_8_5* | 4.94E-05 | 1.38E-06 | 9.82E-05 | 4.30 |
| *m_8_6* | 5.72E-05 | 1.73E-06 | 9.69E-05 | 4.23 |
| *m_8_7* | 5.66E-05 | 1.31E-06 | 9.23E-05 | 4.20 |
| *m_8_9* | 3.43E-05 | 1.31E-06 | 8.52E-05 | 4.19 |
| *m_9_1* | 5.99E-05 | 1.38E-06 | 9.97E-05 | 4.14 |
| *m_9_10* | 4.71E-05 | 1.33E-06 | 8.79E-05 | 3.99 |
| *m_9_11* | 7.21E-05 | 1.58E-06 | 9.64E-05 | 3.75 |
| *m_9_12* | 6.78E-05 | 1.37E-06 | 9.89E-05 | 4.12 |
| *m_9_2* | 7.77E-05 | 2.63E-06 | 9.38E-05 | 4.05 |
| *m_9_3* | 6.47E-05 | 1.91E-06 | 9.63E-05 | 4.19 |
| *m_9_4* | 5.82E-05 | 1.55E-06 | 9.96E-05 | 4.32 |
| *m_9_5* | 5.14E-05 | 1.55E-06 | 9.77E-05 | 4.46 |
| *m_9_6* | 3.93E-05 | 1.41E-06 | 8.75E-05 | 4.54 |
| *m_9_7* | 5.99E-05 | 1.40E-06 | 9.53E-05 | 4.33 |
| *m_9_8* | 3.83E-05 | 1.68E-06 | 9.66E-05 | 3.82 |

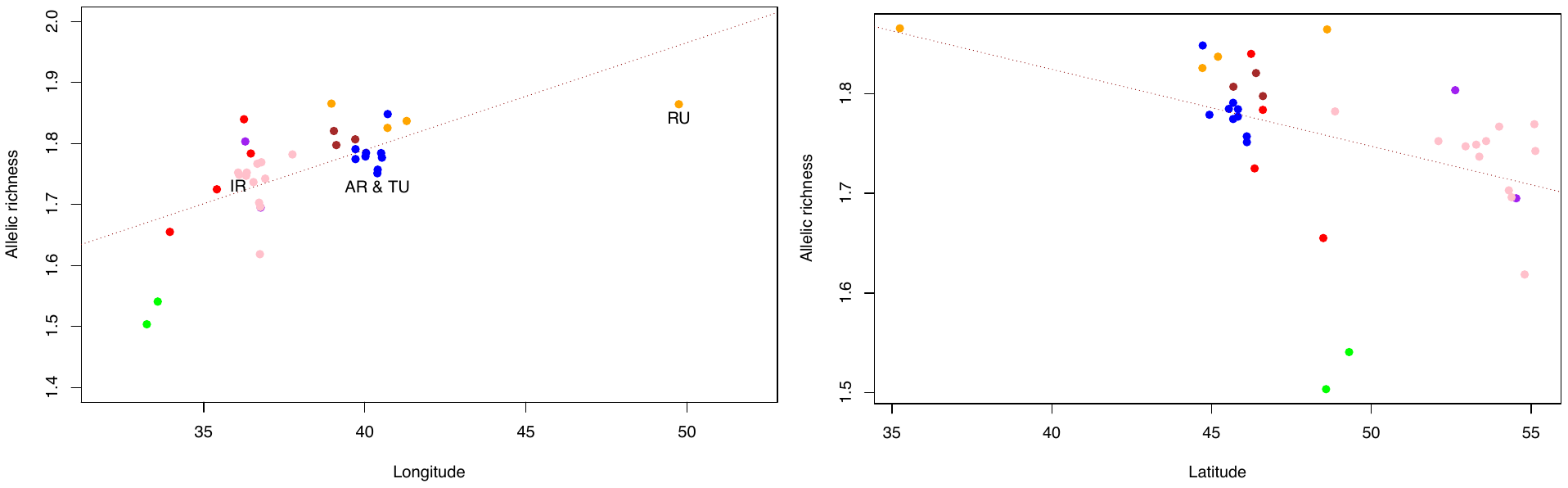

**Figure S11. Spatial genetic variation for *Malus orientalis* (*N*=339, 36 sample sites) in the Caucasus.** Relationships between allelic richness and longitude (left), or latitude (right). Each dot, representing a sampling site, is coloured according to its respective assigned population inferred with STRUCTURE at *K=*9. The red dotted line represents the linear regression line: the adjusted *R^2^* and associated p-value are presented in the main text.

**Table S15.** AUC (ROC curve) and TSS indices for bioclimatic variables for the modeling algorithms (ANN, GAM, GLM) and repetitions for *Malus orientalis.*

|  | Statistic | Repetitions | | | | | Mean |
| --- | --- | --- | --- | --- | --- | --- | --- |
| Models | ANN | 0.83 | 0.70 | 0.84 | 0.70 | 0.80 | 0.77 |
|  | GAM | 0.85 | 0.85 | 0.75 | 0.75 | 0.73 | 0.79 |
|  | GLM | 0.80 | 0.84 | 0.87 | 0.88 | 0.89 | 0.86 |
|  | **Mean ROC** | **0.82** | **0.80** | **0.82** | **0.78** | **0.81** | **0.81** |
| Models | ANN | 0.59 | 0.44 | 0.59 | 0.47 | 0.59 | 0.54 |
|  | GAM | 0.71 | 0.71 | 0.53 | 0.53 | 0.47 | 0.59 |
|  | GLM | 0.59 | 0.65 | 0.68 | 0.71 | 0.76 | 0.68 |
|  | **Mean TSS** | **0.63** | **0.60** | **0.60** | **0.57** | **0.61** | **0.60** |
